# A differential requirement for ciliary transition zone proteins in human and mouse neural progenitor fate specification

**DOI:** 10.1101/2024.02.28.582477

**Authors:** Antonia Wiegering, Isabelle Anselme, Ludovica Brunetti, Laura Metayer-Derout, Damelys Calderon, Sophie Thomas, Stéphane Nedelec, Alexis Eschstruth, Valentina Serpieri, Martin Catala, Christophe Antoniewski, Sylvie Schneider-Maunoury, Aline Stedman

**Author notes:** co-last authors (Sylvie Schneider-Maunoury, Aline Stedman).

## Abstract

Studying developmental processes in the context of the human central nervous system is essential to understand neurodevelopmental diseases. In this paper we perform a comparative functional study of the ciliopathy gene *RPGRIP1L* in human and mouse spinal development using *in vitro* 3D differentiation of pluripotent stem cells. *RPGRIP1L*, a causal gene of severe neurodevelopmental ciliopathies such as Joubert and Meckel syndromes, encodes a scaffolding protein of the ciliary transition zone involved in ciliary gating. Previous work has identified a major role for *Rpgrip1l* in mouse brain and spinal cord development, via controlling the Sonic Hedgehog (SHH)/GLI pathway. We show that spinal organoids derived from *Rpgrip1l* mutant mouse embryonic stem cells faithfully recapitulate the loss of motoneurons and the strong reduction of SHH signaling observed in the mutant mice. In contrast, human induced pluripotent stem cells mutant for *RPGRIP1L* produce motoneurons and activate the SHH pathway at levels similar to wild types, a property shared by human iPSCs mutant for another ciliopathy gene *TMEM67*. Moreover, we show that, in human *RPGRIP1L* mutant organoids, motoneurons acquire a more anterior identity, expressing *HOX* genes and other proteins normally present in the hindbrain and cervical spinal cord while motoneurons from wild type organoids strictly display a caudal brachial identity. By performing a temporal transcriptome analysis throughout the differentiation process, we find that the antero-posterior specification defect arises in early axial progenitors and correlates with the loss of cilia in these cells. Thus, this study uncovers distinct functions in humans and mice for ciliopathy proteins and a novel role for RPGRIP1L in human spinal antero-posterior patterning. These findings have important implications for understanding the role of cilia in human spinal cord development and the pathogenic mechanisms of neurodevelopmental ciliopathies.

## Introduction

Primary cilia are highly conserved signaling organelles that protrude from the surface of most vertebrate cells. Constructed from a microtubule scaffold, the ciliary axoneme is anchored in the plasma membrane via the basal body, a modified mother centriole. A specific structure above the basal body, the ciliary transition zone (TZ), regulates the entry and exit of proteins, thereby acting as a “ciliary gate” that regulates the composition of cilia and creates an independent cell compartment. By modulating several signaling pathways such as SHH, WNT and PDGFR, cilia play essential roles in embryonic development and adult tissue homeostasis. Consequently, defects in primary cilia can lead to a large group of rare genetic diseases called ciliopathies^1, 2^, which can affect the development of multiple organs, particularly the retina, kidneys, lungs, skeleton, heart and the central nervous system (CNS). Ciliopathies are characterized by considerable phenotypic and genotypic overlap and, rather than being considered separate clinical entities, they are now recognized as a spectrum of genetic diseases. Additionally, there is extensive phenotypic and genetic heterogeneity amongst ciliopathies, and mutations in the same gene can cause strikingly different phenotypes, without necessarily any genotype-phenotype correlation.

Although this is not completely deterministic, mutations in genes encoding ciliary TZ proteins often lead to neurodevelopmental ciliopathies such as Joubert Syndrome (JS) and the fetal lethal Meckel Gruber syndrome (MKS)^1, 3^. Most of the current knowledge about neurodevelopmental defects in ciliopathy patients and their underlying mechanisms was gained by *in vivo* studies on animal models and *in vitro* analyses. Using these approaches, it was demonstrated that primary cilia are present throughout the neuronal lineage, from neural stem cells to progenitor cells, mature neurons and glia, where they play an essential role in CNS formation by modulating signaling pathways and cell cycle progression^4–10^. We and others have demonstrated that the ciliopathy causal gene *RPGRIP1L* encodes a large scaffolding protein acting as a keystone of TZ construction^11–19^. In addition, it regulates proteasomal activity at the ciliary base and autophagy in murine cells^20, 21^ and is involved in planar cell polarity in zebrafish and mice^22^. Furthermore, an essential function for Rpgrip1l in SHH signaling was demonstrated in mouse embryonic tissues and cultured cells^11, 15, 16, 23^. However, our understanding of the role of RPGRIP1L and other ciliary proteins in human neural fate is still very partial and has been hampered due to the paucity of human models, limiting our ability to understand the role of cilia in human development, and the origin of neural developmental defects in ciliopathies. Studies using increasingly diverse hiPSC-based organoid models in which the role of ciliary proteins can be investigated are therefore of growing interest to the field. These include studies that investigate ciliary proteins in retinal^24^, cortical^25, 26^ or renal^27^ organoids.

Moreover, evidence suggests that primary cilia function could be only partially overlapping between human and animal models, partly due to specificities in human CNS development. For example, while *Rpgrip1l* mutant mouse embryos, like other ciliary mutants, display a severe disorganization of the ventral diencephalic and hypothalamic regions and of the ventral spinal cord, no such phenotype was described in ciliopathy patients^6, 7, 28–30^.

In this study, we took advantage of pluripotent stem cell-based organoid approaches to study the role of the TZ proteins RPGRIP1L and TMEM67 in human neural progenitor (NP) fate. Both proteins are involved in the gating function of the TZ and have been shown to regulate SHH-dependent spinal patterning in mouse models ^8, 31–33^. However, they are structurally and functionally different: RPGRIP1L is a scaffolding protein and a cornerstone of TZ assembly^18, 19^, while TMEM67 is a Frizzled-like transmembrane receptor^8, 31^, and the two proteins belong to different TZ protein complexes^34^. Since the role for ciliary proteins in SHH-dependent patterning of the ventral spinal cord has been largely studied^35^, we used wild-type (WT) and Rpgrip1l-deficient mouse embryonic stem cell (mESC)- and human induced pluripotent stem cell (hiPSC)-derived spinal organoids to compare the functions of RPGRIP1L in ciliogenesis and spinal progenitors between these two models. We show that mouse *Rpgrip1l^-/-^* organoids exhibit altered dorso-ventral patterning in response to SHH, similar to mutant mouse embryos, emphasizing the cell-intrinsic requirement for Rpgrip1l in spinal progenitors for cilia-dependent SHH pathway activation and commitment into ventral neuronal cell types. In contrast, RPGRIP1L-deficient human iPSCs submitted to a similar differentiation protocol are still able to form primary cilia and adopt SHH-dependent ventral spinal fates. Interestingly, this phenotype is not unique to RPGRIP1L deficiency, but is also observed in TMEM67-deficient human ventral organoids. Remarkably, in addition to the species-specific differences of TZ proteins in cilia formation and signaling between mouse and human, we discovered a novel role for the human RPGRIP1L protein in the precise assignment of antero-posterior (rostro-caudal) identities of human spinal progenitors. Indeed, mutant motoneurons (MNs) derived from early neuromesodermal progenitors (NMPs) shift their identity from thoracic to more rostral/hindbrain identities. This is accompanied by a temporary loss of cilia in early RPGRIP1L-deficient axial progenitor cells. With this study, we bring a first proof of concept that inter-species differences can be dissected by dynamically comparing mouse and human *ex vivo* organoids and we highlight new human-specific functions for the ciliopathy gene *RPGRIP1L* in antero-posterior patterning of axial progenitors.

## Results

### Defective SHH-induced motoneuron specification in *Rpgrip1l* KO mouse spinal organoids

In vertebrates, graded SHH signaling along the dorso-ventral (DV) axis of the spinal cord is necessary for the specification of the floorplate (FP) and the four different ventral progenitor domains p3, pMN, p2 and p1^36–41^ (Fig. 1a). The study of mouse ciliary mutant embryos demonstrated that primary cilia are required for SHH transduction and the correct establishment of spinal DV patterning^15, 35, 42, 43^. In *Rpgrip1l^-/-^* (KO) mouse embryos, brain and spinal cord progenitors lack a functional cilium, leading to altered SHH pathway activation and repression^6, 15, 16^. As a consequence, in the spinal cord, the FP and p3 territories are not established, fewer pMNs are specified and the dorsal domains are shifted ventrally^6,15^. In order to test whether this phenotype can be reproduced *ex vivo*, even under conditions where SHH activation is forced by the addition of a constant dose of SHH agonist, we established WT and *Rpgrip1l KO* mESC lines from mouse blastocysts that we differentiated into ventral spinal organoids^44–46^ . Briefly, mESC were differentiated into embryoid bodies (EBs) in N2B27 neuralizing medium for 2 days (d0-d2). EBs were then targeted to an axial progenitor fate by combined exposure to the WNT agonist CHIR99021 and to retinoic acid (RA), and further ventralized by addition of SAG (Smoothened AGonist; SHH activation). By day 4, embryoid bodies had organized into spinal organoids that comprised spinal neuronal progenitors from different ventral identities, in particular from the motoneuron lineage. Organoids were let to differentiate until day 7, when most progenitors underwent differentiation (Fig. 1b). WT and *Rpgrip1l KO* spinal organoids were collected at different time points of differentiation, sectioned and immunostained to assess the identity of the different progenitors obtained. We found that on average, 40% of spinal progenitors from control organoids committed to a pMN fate by day 4 of differentiation, as evidenced by Olig2 staining (Fig. 1c, d). pMNs progressively differentiated into post-mitotic MNs until day 6, a stage when only Islet1-positive MNs could be detected (Fig. 1c). Control organoids were also composed of progenitors from the more ventral domains p3 and FP, since we could detect the presence of [Nkx6.1+, Olig2-], Nkx2.2+ and FoxA2+ cells (Fig. 1d). Moreover, control organoids nicely recapitulated the sequential activation of ventral neural identities observed *in vivo* in the mouse spinal cord, as *Olig2* expression preceded the initiation of *Nkx2.2* and *FoxA2*.

**Figure 1:**
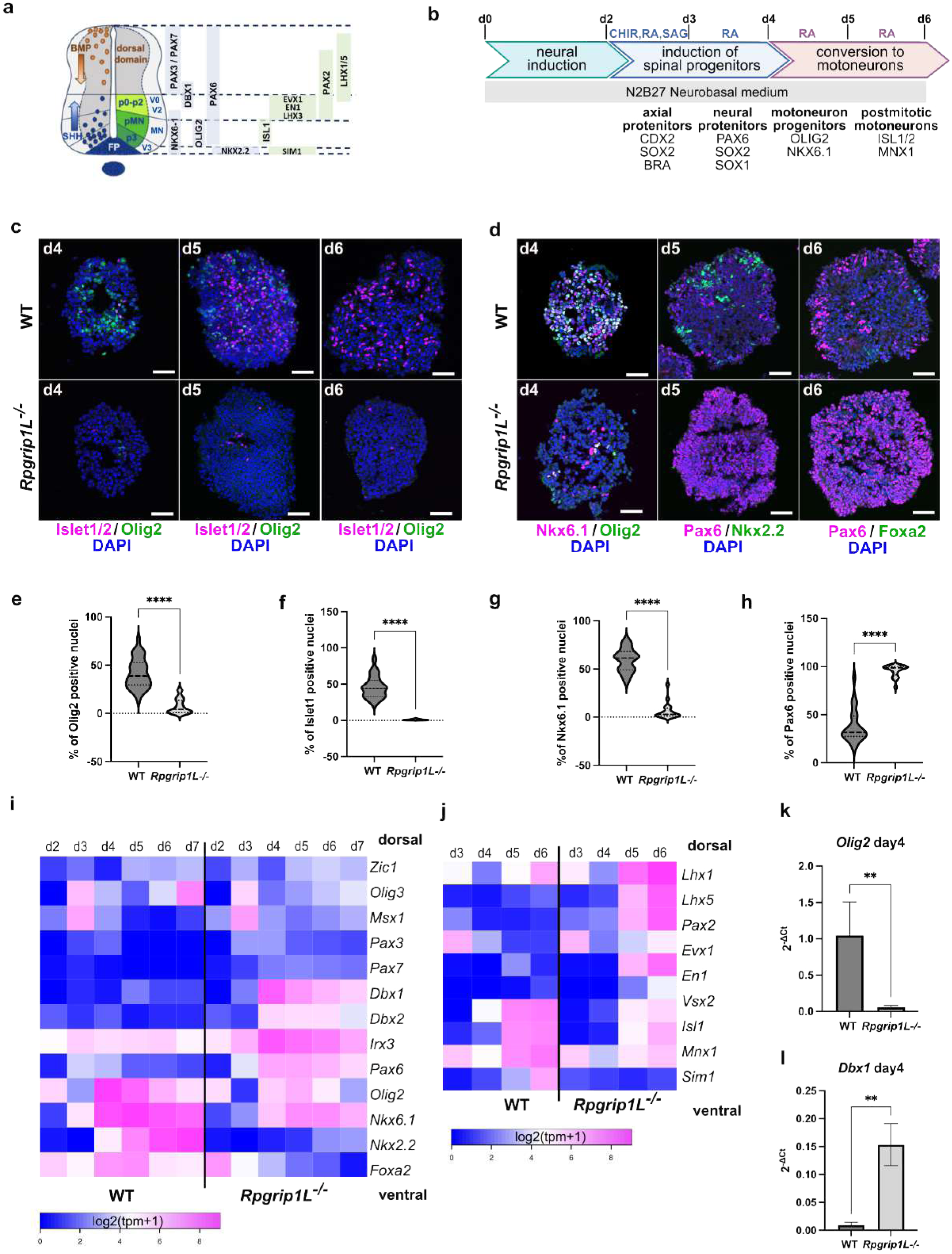
Rpgrip1l-deficient mouse spinal progenitors fail to adopt SHH-dependent ventral fates. **a** Diagram depicting the main transcription factors spanning the different dorso-ventral progenitor (light blue) or neuronal (light green) domains of the developing spinal cord. FP: floor plate. p0-p2, pMN and p3 are ventral spinal progenitor domains. V0-V2, MN, V3 are ventral spinal neuronal domains. **b** Schematic summary of the spinal 3D differentiation approach. The main neural progenitor and neuronal populations obtained at each time point with their molecular markers are indicated at the bottom. Small molecules used to induce or repress signaling pathways are indicated on top. CHIR: WNT agonist CHIR99021; SAG: SHH agonist; RA: Retinoic Acid. Modified after Maury et al., 2015 [56]. **c** Immunofluorescence for Olig2 (pMN) and Islet1 (post-mitotic MNs) in sections from WT and *Rpgrip1l*^-/-^ organoids at indicated time points. d: day of differentiation. **d** Immunofluorescence for Nkx6.1 (p3-pMN-p2), Olig2 (pMN), Nkx2.2 (p3), FoxA2 (FP) and Pax6 (pMN-dp) on sections from WT and *Rpgrip1l*^-/-^ organoids at indicated time points. In **c** and **d**, nuclei are stained with DAPI. Scale bars: 50 μm. **e-h** Violin plots quantifying the percentage of nuclei per organoids stained for the selected markers on sections from WT and *Rpgrip1l*^-/-^ organoids at day 4 (Olig2, Nkx6.1) and day 5 (Islet1, Pax6) of the differentiation. The median and quartiles are indicated by dotted lines. ****P < 0.0001 (unpaired t tests with Welch’s correction). n ≧ 8 organoids analyzed from each genotype, from N≧ 3 experimental replicates. **i, j** Heatmap from bulk RNAseq data based on values of log2(tpm+1) (normalized counts) for selected genes in WT and *Rpgrip1l KO* spinal organoids over time. N=3 experimental replicates for each genotype. **k, l** qPCR analysis for *Olig2* and *Dbx1* in WT and *Rpgrip1l*^-/-^ organoids at day 4. N=5 experimental replicates. Data shown as mean ± SEM. **P < 0.05 (Mann-Whitney).

In contrast, *Rpgrip1l-*deficient progenitors failed to commit to ventral spinal identities. Indeed, the percentage of Olig2+ and Islet1+ cells of the MN lineage as that of Nkx6.1+ cells, was strongly reduced in mutant organoids (Fig. 1c-g), while Nkx2.2+ (p3) and FoxA2+ (FP) progenitor cells were nearly absent from mutant organoids (Fig. 1d). Instead, 80% of the progenitors were positive for PAX6, indicating a shift towards more dorsal identities (Fig. 1d, h). To gain a deeper understanding of the changes in cell fate occurring in *Rpgrip1l-*deficient organoids, we performed bulk RNAseq on WT and mutant organoids from day 2 to day 7, obtained from 3 independent differentiation experiments (Supplementary Fig. 1a). Our transcriptomic analysis confirmed the shift towards dorsal identities in *Rpgrip1l* mutant organoids, and showed that SHH-dependent ventral cell types were replaced by progenitors expressing markers characteristics of the p0-p2 domains such as *Dbx1* (Fig. 1i and Supplementary Fig. 1b-c). qPCR performed on day 4 organoids confirmed these results and showed a more than 10-fold decrease of *Olig2* expression while *Dbx1* expression was increased 5 times in mutant compared with control organoids (Fig. 1k-l). Consistently, the molecular signature of post-mitotic neurons changed from V3-MNs in the control, to V1-V0 in *Rpgrip1l KO* organoids (Fig. 1j). The temporal analysis of the pan-neural markers *Sox1* and *Tuj1* expression in the time course of the differentiation (Supplementary Fig. 1e-f) showed that the reduction in MNs correlated with a transient decrease in neurogenesis between days 3 and 7 in *Rpgrip1l KO* compared with control organoids. Since MNs are the first neurons to differentiate in the spinal cord, this result is in line with the dorsalization occurring in the mutant organoids. In *Rpgrip1l* mutant mice, the defective patterning of the ventral spinal cord has been attributed to an absence of primary cilia in spinal progenitors, and the consequent defect in SHH pathway activation^6, 15^.

Ensemble Gene Set Enrichment Analysis (EGSEA) performed on differentially expressed genes in day 3 and day 4 organoids showed an enrichment in genes related to spinal patterning (Supplementary Fig. 1c) and SHH signaling (Supplementary Fig. 1d). Consistently, qPCR performed on control and mutant organoids showed that while expression of direct targets of the pathway, *Gli1* and *Patched1* was upregulated in controls from day 2 upon SAG addition into the EB culture medium, their expression barely increased in the mutants, showing impaired activation of SHH transduction in absence of Rpgrip1l (Fig. 2a, b and Supplementary Fig. 1g, h). To test whether this phenotype was correlated with defective ciliogenesis we analyzed the ciliary marker ADP Ribosylation Factor Like GTPase 13b (Arl13b) by immunostaining on organoid sections. We found that, similar to what was observed *in vivo*, Rpgrip1l deficiency led to a drastic reduction of Arl13b+ cilia (Fig. 2c, d). This phenotype might reflect a ciliary gating defect, or an absence of ciliary axoneme. To discriminate between these scenarios, we labeled the axonemal protein Ift88 and measured the length of Ift88 staining to evaluate the axonemal length. We found that the axoneme of mutant progenitors was abnormally stunted, with a mean length of 0.2 µm compared to 1 µm in the control (Fig. 2c, e), suggesting that only the TZ might be present above the basal body. This ciliary phenotype was correlated with the complete absence of Rpgrip1l at the ciliary TZ in mutant organoids (Fig. 2c). Overall, our results demonstrate that, as *in vivo*, Rpgrip1l is required *in vitro* in mouse spinal progenitors for cilia structure, SHH pathway activation and the subsequent acquisition of spinal ventral identities. Previous studies showed that spinal organoids can be applied to dissect the activity of the Hedgehog pathway in NPs^46^. Our results provide a proof of concept that they also provide a relevant model for the study of ciliary functions in neural cell fate.

**Figure 2:**
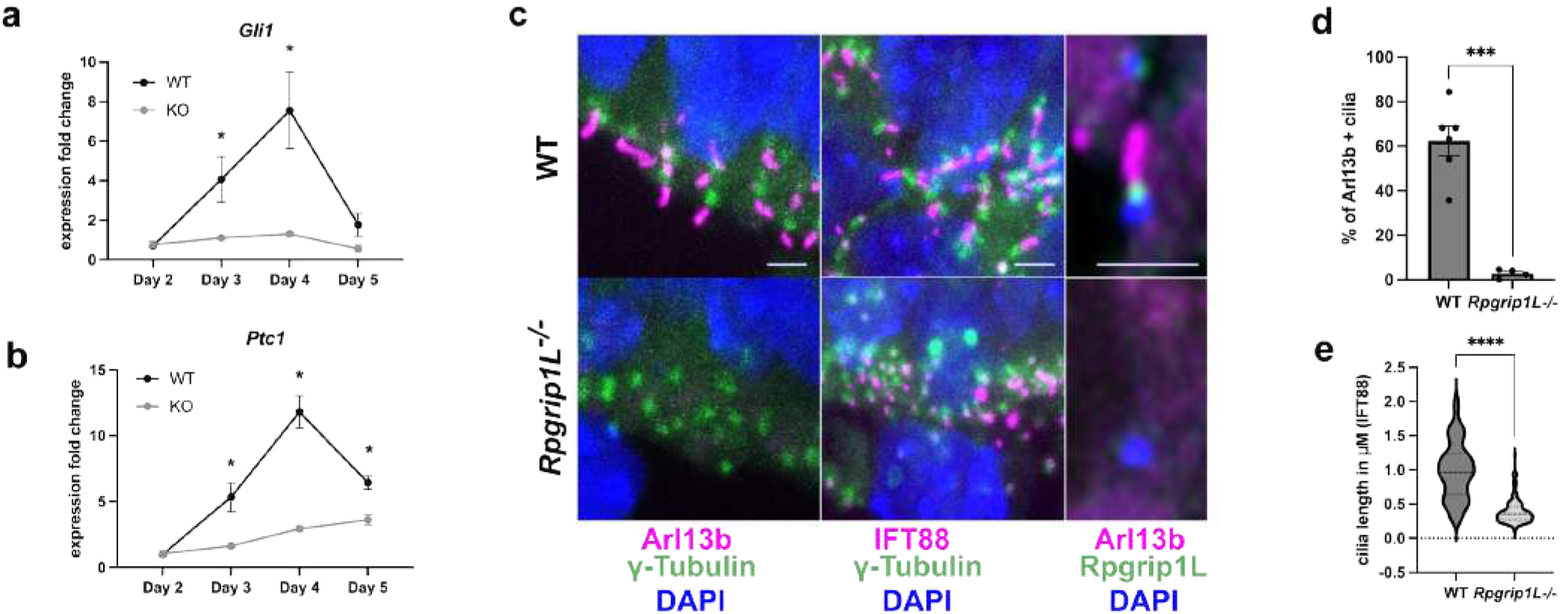
Rpgrip1l-deficient mouse spinal organoids fail to activate the SHH pathway and lack functional cilia. **a, b** Relative fold-change for *Gli1* and *Ptc1* expression in WT and *Rpgrip1l^-/-^* organoids at indicated time points. N=4 experimental replicates. Data shown as mean ± SEM. **P < 0.05 (Mann-Whitney). **c** Immunofluorescence for the indicated ciliary markers on WT and Rpgrip1l-deficient spinal organoids at day 5. Cilia are present at apical sites pointing outside or into inner cavities of the organoids. Scale bar, 2 μm. **d, e** Quantifications of ciliary stainings. Data shown are mean ± SEM. n≧20 cilia from n≧3 organoids were analyzed per genotype. N≧2 independent experiments. ***P < 0.001, ****P < 0.0001 (unpaired t tests with Welch’s correction).

### Human RPGRIP1L-deficient neural progenitors differentiate into MNs in spinal organoids

To tackle the function of primary cilia and the TZ protein RPGRIP1L in human spinal development and SHH transduction, we generated human iPSC clones deficient for RPGRIP1L (*RPGRIP1L^-/-^*) via CRISPR/Cas9-mediated genome editing that lead to premature stop codons in two independent hiPSC lines, PCIi033-A and UCSFi001-A (Supplementary Fig. 2a, b). After genome editing, two *RPGRIP1L*^+/+^ (WT) and two *RPGRIP1L^-/-^* (KO) clones of each hiPSC line (WT: n=4; KO: n=4) were validated (Supplementary Fig. 2b-e) and used for differentiation into ventral spinal organoids following a well-established 3D differentiation protocol^47, 48^ (Fig. 3a). Starting the differentiation of ventral spinal organoids from single cell hiPSCs, EBs were formed after one day in a floating culture in neural induction medium (dual SMAD inhibition) containing the GSK-3β antagonist CHIR99021 for WNT activation and axial progenitor specification^47, 48^ (Fig. 3a). From day 4 onwards, axial progenitors were treated with RA and high concentrations of SAG in order to induce a ventral neural fate (Fig. 3a). At day 6 of differentiation, the majority of WT and RPGRIP1L-deficient cells expressed SOX2 and PAX6, both markers of early NPs, indicating that control cells as well as RPGRIP1L-deficient cells adopted a neural fate (Supplementary Fig. 3a). In response to SAG stimulation, the pMN marker OLIG2 started to be expressed between day 4 and day 6 of differentiation with an expression peak reached at day 9 where 75% of the cells within organoids expressed OLIG2 (Fig. 3b and Supplementary Fig. 3b). Combined to that, cells expressed the broad ventral marker NKX6.1 at day 9, further evidence of ventral spinal fate specification (Supplementary Fig. 3b). Surprisingly, no difference in OLIG2 and NKX6.1 pattern was observed between control and RPGRIP1L-deficient organoids (Fig. 3b and Supplementary Fig. 3b). In line with this, control and RPGRIP1L-deficient cells started to express *ISLET1* and *ISLET2*, markers for postmitotic MNs, from day 9 onwards (Supplementary Fig. 3c-e). A final treatment of organoids with the γ-Secretase Inhibitor DAPT was used to inhibit NOTCH signaling and accelerate the neural differentiation into postmitotic MNs from day 9 onwards (Fig. 3a). Consequently, *ISLET* RNA and protein levels were strongly upregulated in WT and RPGRIP1L-deficient organoids between day 11 and day 14 (Fig. 3b and Supplementary Fig. 3c-f). Taken together, these results show that RPGRIP1L-deficient hiPSCs are able to process differentiation from stem cells to NPs and further from pMNs to postmitotic MNs. Nevertheless, our data reveal 15% less ISLET+ MNs at the end of differentiation in RPGRIP1L-deficient hiPSC-derived spinal organoids compared to controls (Fig. 3b and Supplementary Fig. 3f).

**Figure 3:**
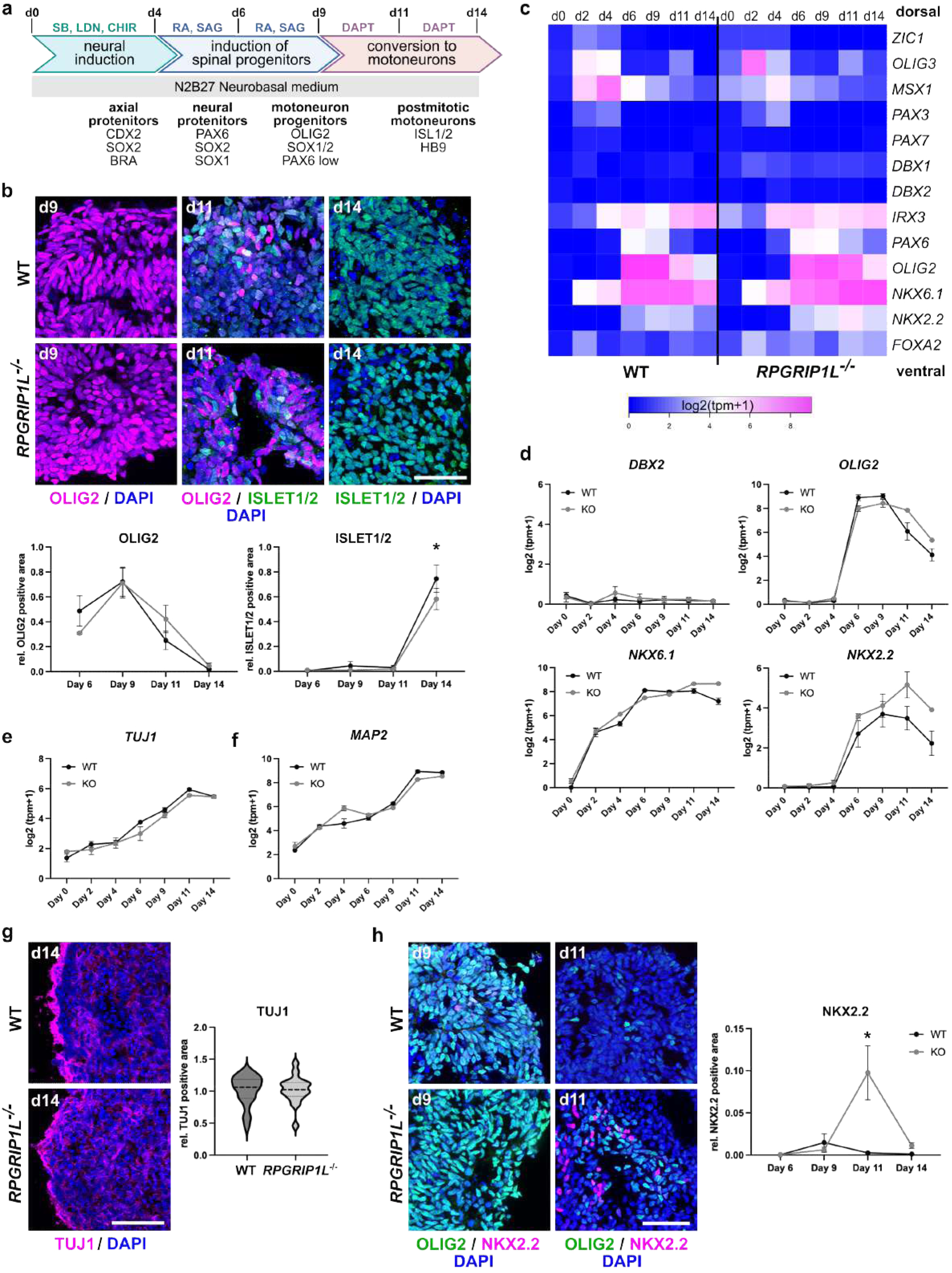
RPGRIP1L-deficient hiPSC-derived spinal organoids adopt the MN fate. **a** Schematic summary of the spinal 3D differentiation approach. Main neural progenitor and neuron populations obtained at each time point with their molecular markers are indicated at the bottom. Drugs used to induce or repress signaling pathways are indicated on top. SB and LDN: SMAD antagonists SB-431542 and LDN-193189; CHIR: WNT agonist CHIR99021; SAG: SHH agonist; DAPT: NOTCH agonist; RA: Retinoic Acid. Modified after Maury et al., 2015 [56]. **b** Immunofluorescence analysis of pMN (OLIG2) and MN (ISLET1/2) production in WT and *RPGRIP1L*^-/-^ spinal organoids. Scale bar: 50 µm. Quantifications show relative OLIG2 and ISLET1/2 positive areas per organoid over time. Data are shown as mean ± SEM. Asterisks denote statistical significance according to unpaired t tests with Welch’s correction (*P < 0.05) (WT: n=4; KO: n=4; N≧3). **c** Heatmap of selected gene expressions from bulk RNAseq data based on log2(tpm+1) values of WT and RPGRIP1L-deficient spinal organoids over time (WT: n=4; KO: n=2). **d** Log2(tpm+1)-graphs illustrate the dynamic expression levels of *DBX2*, *OLIG2*, *NKX6.1* and *NKX2.2* in WT and *RPGRIP1L*^-/-^ organoids. Data are shown as mean ± SEM (WT: n=4; KO: n=2). **e, f** Log2(tpm+1) data from bulk RNASeq analyses show *TUJ1* (**e**) and *MAP2* (**f**) expression in WT and RPGRIP1L-deficient organoids during differentiation. Data are shown as mean ± SEM (WT: n=4; KO: n=2). **g** Immunofluorescence analysis of TUJ1 on day 14. Scale bar: 100 µm. Data are shown as median with quartiles (WT: n=4; KO: n=4; N≧3). **h** Immunofluorescence analysis of OLIG2-positive MN progenitors and NKX2.2-positive spinal progenitors at day 9 and day 11. Scale bar: 50 µm. Data are shown as mean ± SEM. Asterisks denote statistical significance according to unpaired t tests with Welch’s correction (*P < 0.05) (WT: n=4; KO: n=4; N≧3).

By analyzing the expression of various transcription factors that are distinctly expressed in specific DV regions of the developing spinal cord using bulk RNAseq data, we tested whether, despite the normal pMN production, human iPSC-derived organoids deficient for RPGRIP1L exhibited a shift in DV patterning towards the dorsal domains, as it was demonstrated for mESC-derived *Rpgrip1l KO* spinal organoids (Fig. 1). The unaltered expression of the broad ventral marker *NKX6.1*, the intermediate markers *DBX1* and *DBX2*, and the dorsal spinal markers *PAX3*, *PAX7*, and *ZIC1* (Fig. 3c, d) during differentiation, indicated that, contrary to mouse Rpgrip1l-deficient spinal organoids, human RPGRIP1L-deficient spinal organoids did not display a more dorsal progenitor type specification. We noted minor changes among the markers *MSX1* and *OLIG3* at day 2 and day 4 in *RPGRIP1L KO* organoids (Fig. 3c), likely reflecting subtle differences in axial progenitor, before dorso-ventral neural specification.

To investigate whether the mildly reduced number of MNs in RPGRIP1L-deficient organoids could be caused by a delayed neurogenesis, we analyzed the expression of the pan-neuronal markers *TUJ1* and *MAP2*. As expected, expression of both markers increased during differentiation with the highest expression of *TUJ1* and *MAP2* between day 11 and day 14 (Fig. 3e, f) and high protein levels of TUJ1 at day 14 (Fig. 3g). Comparing WT and RPGRIP1L-deficient organoids, no significant differences between *MAP2* and *TUJ1* expression on RNA or protein level was observed. In conclusion, neurogenesis appears unaltered and seems not to be responsible for the slightly reduced number of MNs observed in RPGRIP1L-deficient spinal organoids. Surprisingly, *NKX2.2* RNA and protein expression was significantly increased at day 11 of MN differentiation in RPGRIP1L-deficient cells (Fig. 3d, h). Since NKX2.2 is expressed in various neural progenitors during differentiation, its increased expression could be caused by generation of different cell types in RPGRIP1L-deficient organoids. For instance, NKX2.2 is expressed in visceral pMNs of the hindbrain that do not express OLIG2^49, 50^ as well as in a specific subpopulation of ventral spinal pMNs in humans, where it is co-expressed with OLIG2^51, 52^. Furthermore, NKX2.2 is expressed in the most ventral NPs (p3) and neuronal subtype (V3) within the neural tube (Fig. 1a), where its expression plays a primary role in ventral neuronal patterning and requires high SHH signaling^53^. To test different hypotheses, we first performed co-labeling of OLIG2 and NKX2.2 at day 9 and day 11 of differentiation. We did not detect any increase in OLIG2+/NKX2.2+ cells in RPGRIP1L-deficient organoids compared to WT controls (Fig. 3h and Supplementary Fig. 3g). Consequently, the increased NKX2.2 amount in RPGRIP1L-deficient spinal organoids is unlikely to be caused by increased numbers of ventral and human-specific MN subtypes. However, this does not rule out the possibility of an increased production of hindbrain pMNs in RPGRIP1L-deficient spinal organoids. Next, to test whether increased levels of NXK2.2 could be related to the formation of more ventral NPs and spinal cord neurons, we determined SHH pathway activity in WT and RPGRIP1L-deficient spinal organoids by analyzing the expression profiles of SHH target genes *GLI1* and *PTCH1* along spinal differentiation.

### RPGRIP1L deficient spinal neural progenitor possess cilia and transduce SHH signaling

In both WT and RPGRIP1L-deficient spinal organoids, an increase in *GLI1* and *PTCH1* expression in response to SAG stimulation was observed between day 4 and day 9 of differentiation, followed by an expected decrease in their expression in agreement with the removal of SAG from the differentiation medium (Fig. 4a and Supplementary Fig. 3h, i). No significant difference in SHH target gene expression has been observed between WT and RPGRIP1L-deficient organoids. This result is in line with the functional differentiation of RPGRIP1L-deficient hiPSCs into OLIG2+ pMNs and ISLET+ postmitotic MNs that highly depends on SHH signaling transduction. In contrast, the earlier described increase in the number of NKX2.2+ cells in RPGRIP1L-deficient spinal organoids (Fig. 3d, h and Supplementary Fig. 3g) cannot be explained by an increased SHH pathway activity and a ventralization effect of NPs.

**Figure 4:**
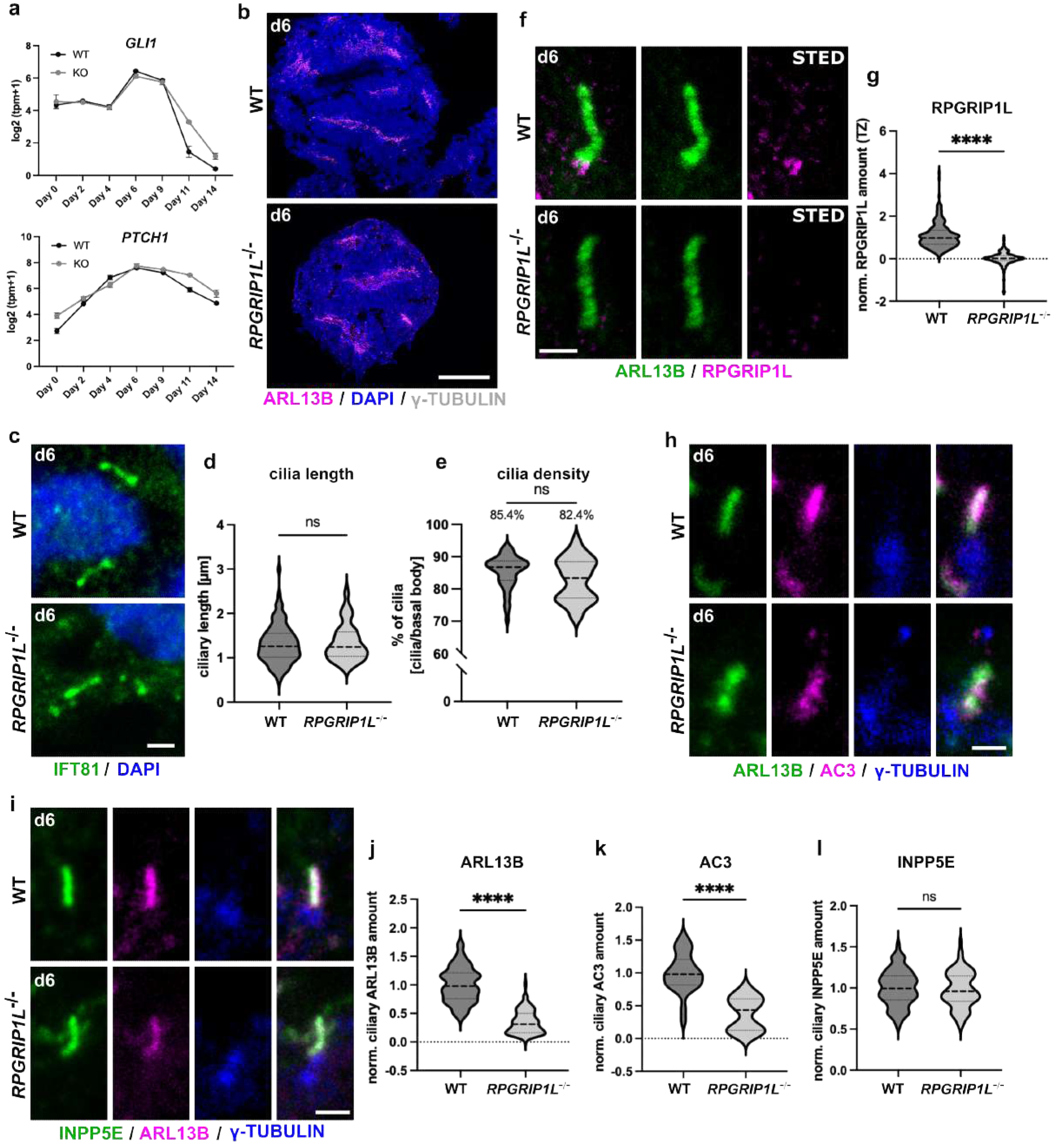
RPGRIP1L-deficient hiPSC-derived neural progenitors harbor cilia and transduce SHH signaling. **a** Log2(tpm+1) data from bulk RNASeq analysis show expression profiles of *GLI1* and *PTCH1* as mean ± SEM (WT: n=4; KO: n=2). **b, c** Immunofluorescence analysis of ciliary proteins in WT and RPGRIP1L-deficient spinal organoids at day 6. **b** Cilia are labeled by ARL13B and basal bodies by γ-TUBULIN. Scale bar: 300 µm. **c** Cilia are labeled by IFT81 in green. Scale bar: 0.5 µm. **d, e** Cilia length and density measurements in WT and *RGPRIP1L^-/-^*organoids at day 6. Data are shown as median with quartiles (WT: n=4; KO: n=4; N≧3). **f** Stimulated-Emission-Depletion (STED) images of WT and RPGRIP1L-deficient cilia at day 6. Cilia are labeled by ARL13B and RPGRIP1L. Scale bar: 0.5 µm. **g** Quantification of the ciliary RPGRIP1L amount based on confocal images. Data are shown as median with quartiles. Asterisks denote statistical significance according to unpaired t tests with Welch’s correction (****P < 0.0001) (WT: n=4; KO: n=4; N≧3). **h, i** Immunofluorescence of WT and RPGRIP1L-deficient cilia on spinal organoids at day 6. Cilia are labeled by ARL13B and AC3 (**h**) or by ARL13B and INPP5E (**i**). Basal bodies are labeled by γ-TUBULIN. Scale bar: 1 µm. **j, k, l** Quantifications of ciliary ARL13B (**j**), AC3 (**k**) and INPP5E (**l**) amounts in WT and RPGRIP1L-deficient organoids. Data are shown as median with quartiles. Asterisks denote statistical significance according to unpaired t tests with Welch’s correction (***P < 0.001) (WT: n=4; KO: n=4; N≧3).

The remaining hypothesis that could explain the increased amount of NKX2.2 in RPGRIP1L-deficient spinal organoids is therefore the production of visceral NKX2.2+ pMNs.

Based on the results of functional SHH transduction and MN differentiation in human RPGRIP1L-deficient spinal organoids, we expected RPGRIP1L-deficient spinal organoids to possess cilia. We analyzed cilia formation at day 6 of MN differentiation, a time point where organoids were stimulated with SAG in order to activate SHH signaling and started to adopt the pMN fate. In WT organoids, short primary cilia were detectable on cells lining the inner lumens of the organoids, thereby pointing into internal cavities (Fig. 4b). These cilia were IFT81-positive and displayed a mean length of 1.3 µm (Fig. 4c, d). RPGRIP1L was present at the ciliary TZ in WT but not RPGRIP1L-deficient spinal organoids (Fig. 4f, g and Supplementary Fig. 2c). Despite the loss of RPGRIP1L, cilia were present lining apical surfaces (Fig. 4b) and displaying an unaltered length and density (Fig. 4d, f) in RPGRIP1L-deficient organoids. However, while WT cilia accumulated the ciliary membrane markers ARL13B, Inositol Polyphosphate-5-Phosphatase E (INPP5E) and Adenylate Cyclase 3 (AC3) (Fig. 4h, i), reduced amounts of ARL13B and AC3 (Fig. 4j, k) but not INPP5E (Fig. 4l) were observed in RPGRIP1L-deficient cilia. This indicates a disrupted ciliary gating in human spinal NPs upon RPGRIP1L deficiency. These results are in line with data from *C.elegans*, different mouse tissues and human cell lines, in which a disrupted TZ organization and in consequence an altered ciliary gating has been described under Rpgrip1l deficiency^17–19, 54–56^. Since ciliary gating can be involved in SHH signaling function^2, 54, 57^ e.g., by regulating ciliary shuttling of pathway components such as SMO (Smoothened) thereby regulating ciliary GPR161 (G Protein-Coupled Receptor 161) amounts^58, 59^, we tested whether GPR161-trafficking was disrupted in RPGRIP1L mutant spinal organoids. GPR161 was enriched in the ciliary membrane in SHH pathway-inactive WT and RPGRIP1L-deficient cells (Supplementary Fig. 3j). In WT and RPGRIP1L mutant NPs stimulated with SAG, GPR161 did not accumulate within primary cilia and thus did not inhibit SHH pathway activation (Supplementary Fig. 3j). These results are in perfect agreement with the previously described functional SHH pathway activation in WT and RPGRIP1L-deficient spinal organoids and show that ciliary gating defects did not affect SHH signaling transduction in RPGRIP1L-deficient NPs.

Given the mild phenotypes observed so far during spinal differentiation of *RPGRIP1L*^-/-^ hiPSCs compared to the results obtained in the murine system, we wondered whether mutations could lead to residual RPGRIP1L activity. To validate the phenotype of our *RPGRIP1L KO* clones, we generated an additional RPGRIP1L-deficient hiPSC clone, this time carrying a full deletion of *RPGRIP1L* (Δexon3-exon27) (Supplementary Fig. 4a, b). The phenotype of this RPGRIP1L-deletion clone was consistent with our previous results obtained from clones carrying *InDel* mutations in *RPGRIP1L*. We observed the loss of RPGRIP1L at the TZ of cilia in NPs (Supplementary Fig. 4c, d) with cilia displaying an unaltered length (Supplementary Fig. 4e) and density (Supplementary Fig. 4f). Furthermore, SHH signaling was unaltered as SHH target gene expression of *GLI1* and *PTCH1* was induced as efficiently in the full deletion *RPGRIP1L* clone as in WT controls (Supplementary Fig. 4g, h). *ISLET1* expression was highly upregulated in control and RPGRIP1L-deficient organoids at day 14 (Supplementary Fig. 4i) and quantification of ISLET+ cells at day 14 validated the efficient generation of MNs in control and RPGRIP1L-deletion clones (Supplementary Fig. 4j, k). Taken together, the consistency of the results demonstrates the authenticity of the phenotype of all our *RPGRIP1L* mutant spinal organoids. To test whether the results obtained in *RPGRIP1L KO* spinal organoids could be extended to other neurodevelopmental ciliopathy genes, we generated TMEM67-deficient hiPSCs via CRISPR/Cas9-mediated genome editing in the PCIi033-A hiPSC line. One control (*TMEM67*^+/-^) and one KO (*TMEM67*^-/-^) clone were used for further analyses (Supplementary Fig. 5a-d). Cilia were present on control and TMEM67-deficient organoids at day 6 of differentiation where they are supposed to transduce SHH signaling (Supplementary Fig. 5c, e). No alterations of cilia length or quantity have been observed (Supplementary Fig. 5f, g), nonetheless gating defects might be present indicated by lower ciliary ARL13B intensities (Supplementary Fig. 5c). As observed in RPGRIP1L-deficient NPs, gating defects in TMEM67-deficient organoids did not affect SHH transduction demonstrated by unaltered SHH target gene expression of *GLI1* and *PTCH1* (Supplementary Fig. 5h, i). Furthermore, control and *TMEM67 KO* organoids started to express *ISLET1* between day 9 and day 14 of differentiation (Supplementary Fig. 5j) without significant differences in the percentage of ISLET+ cells per organoid at day 14 (Supplementary Fig. 5k). Thus, human iPSCs mutant for two distinct ciliopathy genes encoding TZ proteins, *RPGRIP1L* and *TMEM67*, are able to form cilia, transduce the SHH pathway and differentiate into MNs, strongly suggesting that human and mouse spinal progenitors react differently to the loss of TZ proteins.

### RPGRIP1L-deficient spinal organoids show antero-posterior patterning defects with MNs adopting hindbrain identities

To extend the analyses of WT and *RPGRIP1L KO* organoids beyond consideration of MN differentiation, we analyzed the bulk RNAseq data in greater detail. Principal Component Analysis (PCA) demonstrated that WT and KO clones clustered closely together at each analyzed time point, indicating low variation between the genotypes (Supplementary Fig. 6a, b). Further testing for overrepresented gene categories via gene ontology (GOSEQ) did not reveal significant changes in biological processes, molecular functions or cellular components between WT and RPGRIP1L-deficient samples at different time points. Among the few genes that were differentially expressed between WT and RPGRIP1L-deficient organoids, expression of *HOX* genes such as *HOXC6* and *HOXA7* were significantly downregulated in RPGRIP1L-deficient organoids at day 11 and day 14 (Supplementary Fig. 6c, d). In addition, EGSEA on GeneSetDB gene ontology (GO) at day 2 revealed significant downregulation of the biological process *anterior/posterior pattern specification (GO:0009952)* involving downregulation of *HOX* genes as well as the *CDX* genes *CDX1*, *CDX2* and *CDX4* (Supplementary Fig. 6e). In addition to the pairwise comparison of WT and *RPGRIP1L KO* gene expressions at each selected time point, we performed *differential temporal profiles analyzes* on our data. These analyses revealed additional genes whose expression varied significantly over time (Supplementary Fig. 6f), and many of which were related to antero-posterior pattern specification, such as *HOXC8* and *HOXC9* (Supplementary Fig. 6f, g).

*HOX* genes are expressed in four clusters along the rostro-caudal axis of bilateral organisms: HOXA, HOXB, HOXC and HOXD. In humans, HOX proteins are encoded by 39 genes whose regionalized expression is realized by their sequential and collinear activation in axial progenitors, which gradually give rise to mesodermal and neuroectodermal structures with rostral and later caudal identity, each expressing a specific HOX code^48, 60–64^. Importantly, upon differentiation *HOXB* genes become restricted to dorsal spinal neurons and are not expressed in MNs, therefore they were not considered in our analyses^62, 65^. By analyzing the dynamic expression pattern of the remaining 3 *HOX* gene clusters over time, we identified early differences in the expression of posterior *HOX* genes in RPGRIP1L-deficient organoids compared to controls. In controls, early axial progenitors at day 2 started to express anterior *HOX* genes such as *HOXA1* followed by activation of later and more posterior *HOX* genes such as *HOXA7* at day 4 (Fig. 5a), as it was previously reported^48^. At later stages, WT organoids expressed anterior as well as more posterior *HOX* genes (Fig. 5a, d and Supplementary Fig. 7a) as *HOXA* genes from *HOXA1* to *HOXA7* (Fig. 5a, b, c and Supplementary Fig. 7b, c) and *HOXC* genes from *HOXC4* to *HOXC8* (Fig. 5d-f and Supplementary Fig. 7d, e). Consistently to the *HOX* gene expression data, MNs within WT organoids adopted a caudal brachial antero-posterior identity at day 14, that could be visualized by immunofluorescence labeling of HOXA7 and HOXC8, markers for brachial MNs^62^ (Fig. 5g, h), whereas more anterior HOX proteins such as HOXA5, a cervical spinal marker^48, 62^, were not expressed (Fig. 5i). During early stages of differentiation, RPGRIP1L-deficient progenitors expressed anterior *HOX* genes comparable to the WT situation while expression of the more posterior *HOX* genes *HOXA7* and *HOXC8* was already reduced at day 4 (Fig. 5a-f). This difference was even more striking at later stages, where the latest most caudal *HOX* genes expressed in RPGRIP1L-deficient MNs were *HOXA5* instead of *HOXA7* (Fig. 5a, b, c and Supplementary Fig. 7b, c) and *HOXC4/HOXC6* instead of *HOXC8* (Fig. 5d, e, f and Supplementary Fig. 7d, e). These results were confirmed by immunofluorescence analyses, which showed that RPGRIP1L-deficient MNs do not express HOXA7 and HOXC8 (Fig. 5g, h), but significantly increased amounts of HOXA5 (Fig. 5i). Taken together, the differences in *HOX* gene expression indicate a shift in antero-posterior patterning in RPGRIP1L-deficient organoids in which MNs adopt more cervical identities than the caudal brachial identity observed in WT organoids (Fig. 5j) [57, 65, 67].

**Figure 5:**
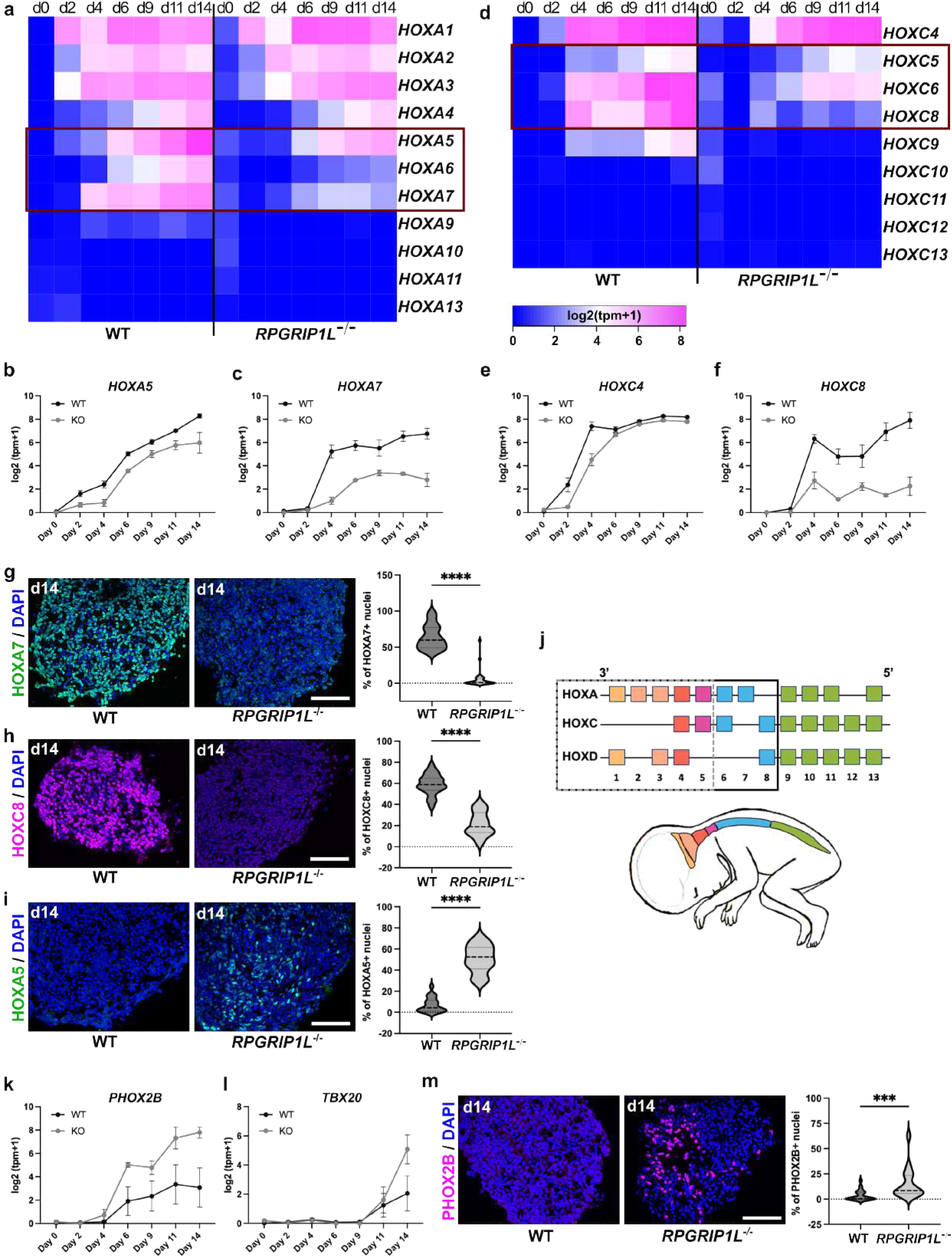
RPGRIP1L-deficient spinal organoids show antero-posterior patterning defects with MNs adopting hindbrain identity. **a, d** Heatmaps showing *HOXA* (**a**) and *HOXC* (**d**) gene expression in WT and RPGRIP1L-deficient spinal organoids over time. The graphs were generated from log2(tpm+1) files of bulk RNASeq analysis (WT: n=4; KO: n=2). **b, c, e, f** Log2(tpm+1) graphs show the expression profiles of selected HOX genes *HOXA5* (**b**), *HOXA7* (**c**), *HOXC4* (**e**) and *HOXC8* (**f**) over time. Data are shown as mean ± SEM (WT: n=4; KO: n=2). **g, h, i** Immunofluorescence of HOX proteins in WT and RPGRIP1L-deficient spinal organoids at day 14. Scale bars: 500 µm. Quantifications show the percentage of HOXA7 (**g**) HOXC8 (**h**) and HOXA5 (**i**) positive nuclei per organoid. Data are shown as median with quartiles. Asterisks denote statistical significance according to unpaired t tests with Welch’s correction (****P < 0.0001) (WT: n=4; KO: n=4; N≧3). **j** Schematic overview about the three HOX gene clusters expressed in MNs from anterior to posterior positions. The black and grey-dotted squares indicate the genes expressed in WT and RPGRIP1L-deficient spinal organoids, respectively. **k, l** Log2(tpm+1) graphs generated from bulk RNASeq analyses show the expression profiles of *PHOX2B* (**k**) and *TBX20* (**l**). Data are shown as mean ± SEM (WT: n=4; KO: n=2). **m** Immunofluorescence of PHOX2B in WT and RPGRIP1L-deficient spinal organoids at day 14. Scale bar: 500 µm. Quantifications show the percentage of PHOX2B positive nuclei per organoid. Data are shown as median with quartiles. Asterisks denote statistical significance according to unpaired t tests with Welch’s correction (*** P < 0.0 01) (WT: n=4; KO: n=4; N≧3).

We further confirmed the anterior shift in MN identities upon loss of RPGRIP1L by analyzing hindbrain specific pMN and MN markers in WT and *RPGRIP1L KO* organoids. Expression of *PHOX2B,* a marker of visceral pMNs and MNs of the hindbrain^66^, and *TBX20*, a late hindbrain-specific MN marker in mice^67^, was increased in *RPGRIP1L KO* compared to WT spinal organoids (Fig. 5k, l and Supplementary Fig. 7f). In line with this, the percentage of PHOX2B+ nuclei per organoid was significantly elevated in RPGRIP1L-deficient organoids (Fig. 5m). The results were confirmed in the full deletion *RPGRIP1L KO* clone (Supplementary Fig. 4l-p), where an increased expression of HOXA5 (Supplementary Fig. 4l) combined with decreased expressions of HOXA7 and HOXC8 was observed (Supplementary Fig. 4l-o). In line with this, the percentage of PHOX2B+ cells per organoid was significantly increased in the full deletion clone of *RPGRIP1L* (Supplementary Fig. 4p).

Interestingly, the results emphasize the previously discussed hypothesis that the increase of NKX2.2+ cells in RPGRIP1L-deficient organoids (Fig. 3d, h) could be explained by the formation of visceral PHOX2B+ MNs.

Analyses of early hindbrain and spine-specific markers further confirmed the shift in antero-posterior patterning upon RPGRIP1L loss. The expressions of the hindbrain patterning markers *EGR2/KROX20*^68–71^, *FGF3*^71–73^, *PHOX2A*^66, 68, 74^ and the visceral pMN marker *NKX2.8*/*NKX2.9*^50, 75^ were strikingly increased in RPGRIP1L-deficient organoids at day 2, day 4 and day 6 (Supplementary Fig. 7g-j). In contrast, expression of *WNT5A*, involved in axis elongation in mammals^76^, was decreased in early *RPGRIP1L KO* progenitors (Supplementary Fig. 7k). In line with this, expression of the spine-specific markers *CDX1* and *CDX4*^77–82^ was strongly reduced in RPGRIP1L-deficient organoids (Supplementary Fig. 7l, m), further emphasizing the caudal to cranial shift in antero-posterior identities of *RPGRIP1L KO* progenitors.

### RPGRIP1L controls axial progenitor fate specification and ciliogenesis in early human spinal organoids

The sequential and collinear expression of *HOX* genes that leads to specification of rostral and later caudal identities is described to be set-up within neuro-mesodermal progenitors (NMPs)^60–63, 83–88^. It has previously been shown that cells in the human spinal organoid model used in this study adopt an axial fate during the first four days of differentiation^47, 48^. This axial fate was defined by the combined expression of CDX2 to BRACHYURY and SOX2, the two main markers of NMPs.

To validate that WT and RPGRIP1L-deficient cells adopted the axial NMP-like fate during differentiation, we analyzed the co-expression of BRACHYURY and SOX2 in organoids at day 2 and day 4. SOX2 was equally expressed in WT and *RPGRIP1L KO* organoids (Supplementary Fig. 8a) with nearly 100% of SOX2+ cells per organoid at day 2 and day 4 (Supplementary Fig. 8c, d, f, g). In contrast, while BRACHYURY expression was equally induced in WT and RPGRIP1L-deficient organoids at day 2 (Supplementary Fig. 8b, c, e), its expression dropped as soon as day 2 in RPGRIP1L-deficient cells (Supplementary Fig. 8f, h) indicating an altered induction of the NMP-like fate upon RPGRIP1L loss. In addition to BRACHYURY and SOX2, axial NMP-like progenitors in WT organoids expressed the caudal marker CDX2 at day 2 and day 4 of differentiation (Fig. 6a-d). Analyses of *CDX2* gene expression profiles showed its strong upregulation between day 0 and day 2 of differentiation, followed by its downregulation from day 4 onwards (Fig. 6e). In RPGRIP1L-deficient organoids, *CDX2* expression was reduced (Fig. 6e) and significantly less CDX2+ cells per organoid were detectable at day 2 and day 4 of differentiation (Fig. 6a-d), with clusters of CDX2-negative cells in RPGRIP1L-deficient but not WT organoids (Fig. 6c). Together, the results of reduced BRACHYURY and CDX2 expression in KO organoids point to a defective induction of the axial NMP-like cell fate upon RPGRIP1L loss.

**Figure 6:**
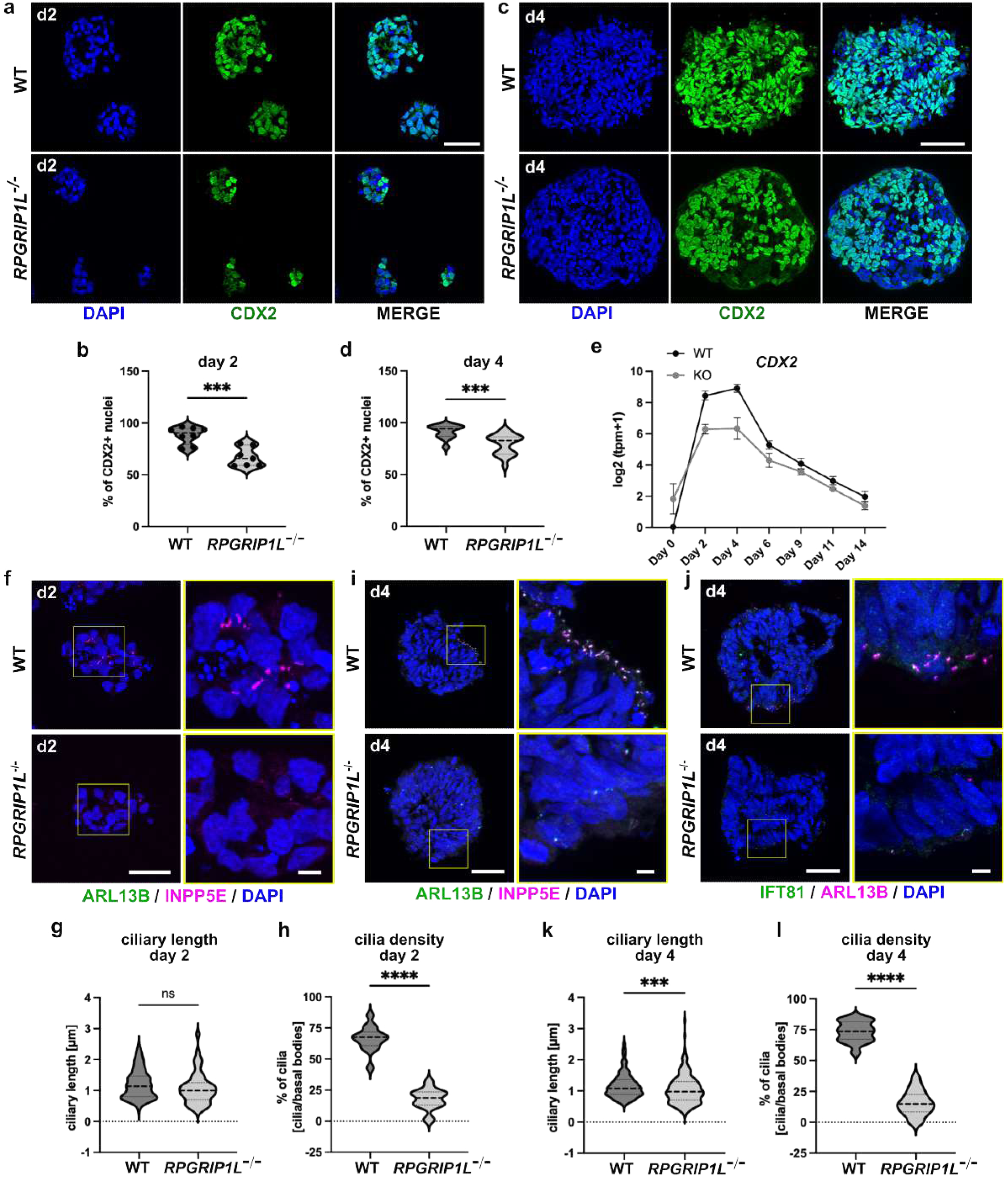
Altered axial patterning and reduced ciliogenesis in early RPGRIP1L-deficient progenitors. **a, c** Immunofluorescence of CDX2 in WT and RPGRIP1L-deficient spinal organoids at day 2 (**a**) and day 4 (**c**). Scale bars: 150 µm. **b, d** Quantifications show the percentage of CDX2 positive nuclei per organoid at day 2 (**b**) and day 4 (**d**). Data are shown as median with quartiles (WT: n=4; KO: n=4; N≧3). Asterisks denote statistical significance according to unpaired t tests with Welch’s correction (***P < 0.001). **e** Log2(tpm+1) graph generated from bulk RNASeq analyses show the dynamic expression profile of *CDX2*. Data are shown as mean ± SEM (WT: n=4; KO: n=2). **f, i, j** Immunofluorescence of cilia in WT and RPGRIP1L-deficient spinal organoids at day 2 (**f**) and day 4 (**i, j**). Cilia are labeled by ARL13B and INPP5E (**f, i**) or by IFT81 and ARL13B (**j**). Scale bars: 150 µm. Magnified areas are indicated by yellow rectangles and magnified images are displayed on the right. Scale bars: 5 µm. **g, h, k, l** Cilia length and cilia density measurements in WT and *RPGRIP1L*^-/-^ organoids at day 2 and day 4. Data are shown as median with quartiles (WT: n=4; KO: n=4). Asterisks denote statistical significance according to unpaired t tests with Welch’s correction (****P < 0.0001).

We next analyzed the ciliary status of axial progenitors at day 2 and day 4 of differentiation. At day 2, cilia were present in WT organoids in an unpolarized fashion (Fig. 6f, g, h) whereas cilia density was significantly reduced in *RPGRIP1L KO* organoids (Fig. 6f, h). The same difference was observed at day 4 of differentiation. Here, cilia were present at the apical sites of WT organoids (Fig. 6i-l), but absent from *RPGRIP1L KO* cells (Fig. 6i-l).

Since the loss of cilia in early RPGRIP1L-deficient organoids correlated with the reduced induction of the axial NMP-like fate, a cell type known to be significantly involved in the establishment of antero-posterior identities, we hypothesize that cilia loss on early RPGRIP1L-deficient progenitors is responsible for the reduced specification of the NMP-like axial fate that lead to the antero-posterior patterning defects observed in *RPGRIP1L KO* organoids. In line with this hypothesis, analyses of TMEM67-deficient organoids revealed that cilia were present on control and KO progenitors at day 4 of differentiation (Supplementary Fig. 9a, b), with *TMEM67 KO* cilia showing the same length and density as controls (Supplementary Fig. 9c, d). Furthermore, the NMP fate was correctly acquired in control and *TMEM67 KO* cells, demonstrated by BRACHYURY and SOX2 expression at day 4 (Supplementary Fig. S9e). Consequently, no antero-posterior patterning defect was observed in TMEM67-deficient organoids at day 14. Expression of the *HOX* genes *HOXA5*, *HOXA7*, *HOXC6* and *HOXC8* was induced between day 4 and day 14 in control and *TMEM67 KO* organoids (Supplementary Fig. 9f) and the percentage of HOXA7+ and HOXC8+ cells per organoid was unaltered in TMEM67-deficient organoids compared to controls (Supplementary Fig. 9g). In both control and KO organoids a few HOXA5+ nuclei were observed (Supplementary Fig. 9g). Furthermore, low expression of *PHOX2B* was detected in control and *TMEM67 KO* organoids at day 14 (Supplementary Fig. 9h), consistent with a few PHOX2B+ cells present in control and KO organoids (Supplementary Fig. 9i).

Interestingly, the correlation between the ciliary phenotype during early stages of differentiation and the axial fate establishment defects seems to be human specific, since no spinal-to-cranial shift of progenitor identity was observed in *Rpgrip1l*^-/-^ mouse organoids, as evidenced by the unaltered expression of *Cdx1-2-4,* and *Cyp26a1* in mutant progenitors compared to controls (Supplementary Fig. 10a-d). No difference in the efficiency of axial progenitors induction was observed in mutant organoids, where at day 3 similarly to control organoids, the majority of cells expressed Cdx2 (Supplementary Fig. 10e-f). Notably, in control and mutant organoids, the same proportion of progenitors co-expressed Brachyury and Sox2 at day 3 suggesting that in both genotypes progenitors transited through a bi-potent NMP-like state (Supplementary Fig. 10g-i). To gain more insight into the antero-posterior identity of control and *Rpgrip1l*^-/-^ mouse spinal progenitors present in our organoids, we also analysed *Hox* gene expression by bulk RNAseq and qPCR analyses. We found that in both genotypes, organoids expressed predominantly brachial and anterior thoracic *Hox* genes (Supplementary Fig. 10i-m). Moreover, although we noticed a slight decrease in *Hox10* gene expression, no major alterations were observed in *Hox* gene expression that might indicate severe antero-posterior specification defects in the mutants (Supplementary Fig. 10i-m). Finally, analysis of cilia in organoids at day 3 revealed that mouse axial progenitors deficient for Rpgrip1l, even though not affected in their axial fate, still presented abnormally shortened cilia (Supplementary Figure 10n-o). Therefore, Rpgrip1l is required in human but not mouse axial progenitors, for early axial identity specification mediated by primary cilia.

## Discussion

Previous analysis of *Rpgrip1l* and *Tmem67* mutant mice demonstrated that deficiency for these TZ proteins led to DV mis-patterning of the neural tube marked by the strong decrease of ventral neuronal subtypes like V3 and MNs^6, 15, 32^. These phenotypes were correlated to the absence of primary cilia and functional SHH signaling, and were also described for *Ift* mutant mice^15, 43, 89^. Since these mice are lacking or displaying a reduced FP, a question remained as to whether altered patterning of the neural tube resulted only from a reduced SHH secretion ventrally, or from impairment of SHH signal transduction within ventral spinal progenitors themselves. Here, by exposing control and *Rpgrip1l* mutant spinal organoids to high dose of SHH agonist, we show that primary cilia are intrinsically required within mouse spinal progenitors to transduce SHH. Upon inactive SHH signaling, mutant progenitors switch on intermediate-dorsal identities instead of ventral ones.

Strikingly, we find that contrary to the mouse, human RPGRIP1L and TMEM67 are not required for the establishment of ventral spinal neuronal cell types in organoids. Despite a clear ciliary gating defect, with decreased ARL13B or AC3 ciliary localization, human mutant spinal NPs were still able to grow primary cilia and transduce SHH signaling similarly to controls, adopting MN identities. Ciliogenesis in the absence of Rpgrip1l and Tmem67 has also been observed in other cellular contexts, like in mouse limbs progenitors, MEFs, HEK cells^18–20, 32^ or human fibroblasts derived from *RPGRIP1L* and *TMEM67* mutated JS patients^17, 90^. Whether the difference in ciliary phenotype observed between mouse and human spinal progenitors stems from differences in ciliary composition or stability, and accounts for the difference in SHH activation in these two systems remains to be addressed. In addition to the species-specificity differences in the ciliary phenotype, our results highlight a context-specific role for ciliary TZ proteins in human neural progenitors. In RPGRIP1L-deficient human organoids, NPs at day 6 display cilia of normal length and a moderately altered ciliary content, while at day 2 and day 4, progenitors display a strong reduction of the cilia density. Differences in ciliary structure and composition between tissues, already suggested by the high heterogeneity of phenotypes in human ciliopathies, have been described in other models^18, 91–93^. However, here we show for the first time a change in cilia sensitivity to the loss of a TZ protein along the process of neural progenitor specification in humans, at a key step of cell fate transition.

Whether this change parallels differences in ciliary structure, and which signaling and molecular cascades lead to these changes in cilia stability during the differentiation process are important questions raised by our study. In this regard, recent data show that cell fate can regulate ciliogenesis by different mechanisms such as transcriptional regulation^94^, niche condition-specific regulation^95, 96^, and several non-canonical signaling pathway activities such as SHH^97^ and WNT^98^. It will be a key future task to examine cell type specific cilia features and explore whether different modes of ciliogenesis regulation play a role in diverse spinal progenitors to explain at least in part the regional specificity of defects found in neurodevelopmental ciliopathies.

Quite unexpectedly, we found that although they do not display SHH-dependent dorsoventral patterning defects, human *RPGRIP1L* mutant spinal organoids shift to more anterior (hindbrain/cervical spinal) identities as assessed by altered HOX expression patterns and activation of genes specific for hindbrain neurons such as *PHOX2B*. This switch occurs at early stages of HOX activation, at the axial progenitor stage (day 2 in our spinal organoids) and is paralleled by the above discussed transient loss of cilia and a reduction in the expression of *CDX* genes. Later (at day 4), reduced numbers of BRACHYURY+/SOX2+ NMPs are found in *RPGRIP1L* mutants. In *TMEM67* mutants, which do not lose cilia at early stages, no perturbation of AP patterning nor NMP-fate establishment is observed; Thus, we find a strong correlation between cilia loss at axial progenitor and NMP stage and the activation of a more anterior progenitor programme in spinal organoids. Why TMEM67 depletion does not lead to cilia loss at day 4, while RPGRIP1L depletion does, remains to be understood. As mentioned earlier, these two TZ proteins have different structures and functions and participate in distinct complexes at the TZ^8, 18, 31, 34^. Thus, it is possible that they contribute differently to TZ stability and ciliary gating. The analysis of other ciliary gene mutants will be essential to understand the contribution of different TZ proteins in cilia stability.

Altered antero-posterior patterning could arise from several mechanisms. WNT, RA, Fibroblast Growth Factor (FGF), and more recently NOTCH signaling, have been involved in the determination of axial progenitors and thereby regulation of gradual *HOX* gene expression^47, 64, 77, 99–103^. From our transcriptome data, we could not observe any indication of perturbations in FGF or NOTCH pathways upon RPGRIP1L deficiency. Analysis of Wnt3a- and Cdx-deficient mouse embryos^78, 104, 105^ or experimental manipulation of CDX or BRACHYURY activity^106–109^ lead to a working model in which early *BRACHYURY* and *CDX* expression activates and fine-tunes *HOX* gene expression profiles along the antero-posterior axis downstream of WNT/β-Catenin signaling in axial progenitors^77, 79, 83, 85, 99, 100, 104, 106, 109–112^. Interestingly, it was demonstrated recently that WNT signaling during early differentiation of hiPSCs into spinal MNs allows progressive, collinear HOX activation and that a switch to RA treatment at any time point during this process stops collinear HOX activation and favors transition into neuroectodermal progenitors with a discrete HOX gene profile^48, 100^. Thus, we hypothesize that the shift in antero-posterior patterning that we describe in RPGRIP1L-deficient human spinal organoids results from reduced WNT transduction during early stages of axial progenitor differentiation, leading to a reduction in BRACHYURY, CDX2, *CDX1* and *CDX4* expression and the consequent decreased expression of posterior *HOX* genes, so that MNs adopt hindbrain or cervical rather than caudal brachial identities. Whether RPGRIP1L or cilia act directly on maintaining WNT signaling in axial progenitors remains to be demonstrated. While cilia and TZ proteins, and specifically RPGRIP1L and TMEM67, have been involved in modulating WNT pathways, their effect is complex and depends on the tissue and cell type^22, 113–117^. Further analyses will show if and how cilia regulate WNT signaling and consequently *BRACHYURY*, *CDX* and *HOX* gene expression during axial progenitor fate specification, or whether other pathways such as FGF and RA are implicated.

Another important question raised by our study is whether the defect in antero-posterior patterning upon loss of RPGRIP1L could be related to phenotypic features of neurodevelopmental ciliopathies. Since the antero-posterior patterning defect of MNs described here most likely lead to a mild shift of boundaries between the posterior hindbrain region and the anterior spinal region with an increase in anterior identities, a corresponding phenotype could be present in ciliopathy patients. Unfortunately, analyses of such patterning defects in patients are not available. A first important step would be therefore to test the differentiation capacity of patient-derived hiPSCs and analyze their phenotype related to antero-posterior patterning. Apart from that, our study provides interesting entry points for further investigations where it will be important to analyze antero-posterior patterning more carefully and in additional CNS regions such as the hindbrain, as it is highly sensitive to antero-posterior patterning defects and one of the major targets of neurodevelopmental ciliopathies. Here, the increased expression of PHOX2B appears to be particularly interesting, since its deregulation was shown to cause breathing pattern regulation defects^118–122^, a feature that can also be observed in JS patients^123^.

## Material and Methods

### Generation of mESCs lines

*Rpgrip1l^-/-^* and *Rpgrip1l^+/+^* mouse ES cell lines were derived from blastocysts according to standard procedures [139] using *Ftm*-deficient mice produced and genotyped as described previously [15], and maintained in the C57BL/6J background. One ES clone from each genotype was used in this study. Both *Rpgrip1l*^-/-^ and *Rpgrip1l*^+/+^ clones are males, which avoids sex-dependent bias in RNAseq data.

### mESCs culture and differentiation into ventral spinal organoids

mESCs were cultured in DMEM-GlutaMAX (Gibco) supplemented with 15% FCS, 100 µM β-mercaptoethanol, 100 U/ml penicillin, 0.1 mg/ml streptomycin, and 10^3^ U/ml LIF (Millipore). To generate mouse spinal organoids, low-passage (inferior to 10) mESCs were dissociated and differentiated into embryoid bodies by seeding them in ultra-low attachment petri dishes (Corning) at a density of 9×10^4^ cells ml^−1^ in N2B27 differentiation medium containing Advanced DMEMF12 and Neurobasal media (1:1, Life Technologies), B27 w/o vitamin A (Life Technologies), N2 (Life Technologies), 2 mM L-glutamine (Life Technologies), 0.1 mM β-mercaptoethanol, penicillin and streptomycin (Life Technologies). On day 2, the medium was changed and supplemented with 3µM CHIR, 10nM RA and 500nM SAG to induce ventral spinal fate. The medium was changed daily, with addition of 10nM RA only.

### hiPSCs culture

hiPSCs were thawed in presence of Rock-inhibitor Y-27632 (5µM, Stemgent-Ozyme #04-0012) and cultured under standard conditions at 37°C in mTeSR+ medium (Stem Cell Technologies #100-0276) on Matrigel (Corning, VWR #354277) coated plates upon confluency of 70-80%. Passaging was performed using ReLeSR (Stem Cell Technologies #05872) and testing for potential mycoplasma contamination was performed regularly by Eurofins genomic (Mycoplasmacheck). Accutase (Stem Cell Technologies **#**07920) was used for the dissociation of hiPSC colonies into single cells.

### Generation of mutant hiPSC lines

*RPGRIP1L* mutant and wildtype hiPSC clones were produced via CRISPR/Cas9-mediated genome editing in two different hiPSC lines: PCIi033-A (PHENOCELL (PLI)) and UCSFi001-A (deposited at the Coriell Institute for Medical Research under the identifier GM25256). Both cell lines are reported to be males. The targeted exon 3 of *RPGRIP1L* and flanking sequences were sequenced prior to gene editing in both cell lines (see *on- and off-target analyses* section). The gRNA targeting exon 3 of *RPGRIP1L* was designed using the online design tool CRISPOR (134). Annealing of crRNA (Alt-R® CRISPR-Cas9 crRNA, 400 nM, IDT) and tracrRNA (Alt-R® CRISPR-Cas9 tracrRNA, 400 nM, IDT #1072533) was performed and 400.000 hiPSCs of each line were transfected with 1µl Cas9 protein (30 µM; kindly provided by TACGENE, Paris, France) and 1µl gRNA (200 µM) using nucleofection. The transfected cells were cultured in mTeSR+ medium (Stem Cell Technologies #100-0276) containing Rock-inhibitor Y-27632 (10 µM, Stemgent-Ozyme #04-0012) for one day, replaced by mTeSR+ medium without Rock-inhibitor. Medium was changed every second day and serial dilutions were performed in mTeSR+ medium containing Rock-inhibitor Y-27632 and CloneR (1:1000, Stem Cell Technologies **#**05889). Rock-inhibitor was removed after two days and the medium containing CloneR was changed every second day until colonies were ready to be picked. Colonies were isolated in 24-well plates in mTesR+ medium containing CloneR. DNA was isolated for genotype analyses (see *on- and off-target analyses* section). Four WT (two clones of each hiPSC line) and four homozygous mutant hiPSC clones (two of each hiPSC line) carrying *InDel* mutations that lead to premature STOP were used for further analyses (Supplementary Fig. 2).

A large deletion of *RPGRIP1L* was induced in the PCli033-A line by combining the gRNA targeting exon 3 of *RPGRIP1L* with a gRNA targeting exon 27 of *RPGRIP1L* (Δexon3-exon27). This approach was performed at the ICV-iPS platform of the Brain Institute in Paris (ICM, Paris) under supervision of Stéphanie Bigou. One *RPGRIP1L KO* clone and one heterozygous control clone were used for further analyses (Supplementary Fig. 3). For the generation of TMEM67-deficient hiPSC clones from the PCli033-A cell line, the following strategy was used: The gRNA targeting exon 1 of *TMEM67* was combined with a gRNA targeting exon 27 of *TMEM67* to generate a large deletion of TMEM67 (Δexon1-exon27). One *TMEM67 KO* clone and one heterozygous control clone were used for further analyses (Supplementary Fig. 5). For all clones, genomic stability was assessed by detection of recurrent genetic abnormalities using the iCS-digitalTM PSC test, provided as a service by Stem Genomics (https://www.stemgenomics.com), as described previously^124^.

### On- and off-target analyses

Genomic DNA was extracted from both hiPSC lines (PCli033-A and UCSFi001-A) prior to CRISPR/Cas9 gene editing. Genotyping of on-targets was performed by PCR, and PCR products were sent for sequencing. After CRISPR/Cas9 gene editing, genomic DNA was isolated from individual hiPSC clones and the on-target sites were amplified by PCR. The PCR products were sent for sequencing to determine the specific mutations that were induced. Off-targets were identified using the CRISPOR online design tool^125^ and ranked according to their CFD off-target score^126^. Potential off-targets within exons and with CFD scores > 0.02^127^ were selected for PCR and subsequent sequencing analyses. The primers used for on- and off-target analyses are listed in Table S1.

### Differentiation of hiPSCs into spinal organoids

Differentiation of wildtype and RPGRIP1L mutant hiPSCs was performed as previously described^47, 48^. After amplification, hiPSC lines were dissociated into single cells using Accutase (Stem Cell Technologies **#**07920) and resuspended in differentiation medium N2B27 [vol:vol; Advanced DMEM/F-12 (Gibco) and Neurobasal Medium (Gibco)] supplemented with N2 (Thermo Fisher #17502048), B27 without Vitamin A (Thermo Fisher #12587010), penicillin/streptomycin 1 % (Thermo Fisher #15140122), β-mercaptoethanol 0.1 % (Thermo Fisher #31350010). Cells were seeded in ultra-low attachment dishes (Corning #3261) to allow EB formation. The N2B27 differentiation medium was used throughout the whole differentiation process, but different additives were given at different time points (Fig. 3a). Rock-inhibitor Y-27632 (5 μM; Stemgent-Ozyme #04-0012) was added from day 0 to day 2, CHIR-99021 (3 µM; Stemgent-Ozyme #04-0004) from day 0 to day 4, SB431542 (20 μM; Stemgent-Ozyme #04-0010) from day 0 to day 3 and LDN 193189 (0.1 μM; Stemgent-Ozyme #04-0074) from day 0 to day 4. The differentiation was preceded by adding Smoothened AGonist (SAG 500 nM; Merck #566660) and Retinoic Acid (100 nM RA; Sigma #R2625) to the N2B27 differentiation medium from day 4 to day 9. γ-Secretase inhibitor DAPT (10 µM; Tocris Bioscience #2634) was added from day 9 to the end of differentiation. Medium was changed every other day and additionally at day 3 of differentiation.

### EB embedding and Cryosectioning

Murine and human spinal organoids were collected at different time points during differentiation, rinsed with PBS and fixed in cold PFA (4 %) for 7-12 min at 4 °C. EBs were rinsed in PBS and incubated in 30 % sucrose in PBS until completely saturated. Cryoprotected EBs were embedded in OCT embedding matrix (Cell Path #KMA-0100-00A) and stored at −80 °C. 12 µM cryostat sections were prepared and stored at −80 °C.

### Immunofluorescence

Cryostat sections of murine and human EBs were washed in PBS/0.1% Triton X-100. Optional steps for permeabilization with PBS/0.5 %Triton X-100 for 10 min, post-fixation with MeOH (100 %) for 10 min or antigen-retrieval with Citrate-Buffer were performed depending on the antibody and specimen. After washing, cryosections were incubated with a blocking solution containing 10 % NGS in PBS/0.1% Triton-X100 for at least 2 hours. The sections were incubated with primary antibodies (Table S2) diluted in PBS/0.1 % Triton-X100 and 1 % NGS overnight at 4 °C. Next day, the slides were washed 3 x 5 min with PBS/0.1 % Triton-X100 and incubated with the secondary antibodies (Table S2) diluted in PBS/0.1 % Triton X-100 and 1 % NGS for 2 hours. After final washing steps with PBS/0.1 %Triton X-100 for at least 1 hour, sections were mounted with Vecta-Shield (Vector #H-1000) or Mowiol (Roth #0713.2) optionally containing DAPI (Merck #1.24653).

### Image acquisition

Fluorescence image acquisition of patterning and differentiation markers was performed at room temperature using a Zeiss Observer Z1 equipped with an Apotome module, an Axiocam 506 monochrome cooled CCD camera and a 40x oil objective with a NA 1.3. All images were processed with Zen software (blue edition). Image analyses and figure preparation were performed using Fiji (ImageJ; National Institutes of Health). For cilia analyses, fluorescence images were acquired at room temperature using an inverted confocal microscope (LSM 710, Carl Zeiss AG or TCS SP5, Leica), a 63x oil objective with a NA 1.4, and a monochrome charge-coupled device camera. Z-stacks with 0.2 µM steps were acquired. Images were processed with Zen software (black edition). Image analyses and figure preparation were performed using Fiji (ImageJ; National Institutes of Health).

STED images were acquired on the STEDYCON confocal and STED module (Abberior Instruments GmbH, Germany) connected to the lateral port of an IX83 inverted microscope (Evident Scientific). The STEDYCON SmartControl software was used for data acquisition. Imaging was performed using a UPLXAPO 100x oil immersion objective (Evident Scientific). For STED, a pulsed 775 nm laser was used. STAR RED was excited with a pulsed 640 nm laser, STAR ORANGE was excited with a pulsed 561 nm laser, and DAPI was excited using a continuous wave 405nm source. Pixel size was 20 nm in xy direction, with a 250 nm step size in the z direction.

### Quantification of fluorescent staining and statistical analyses

The quantitative analysis of nuclear stainings was done either by scoring the percentage of positive nuclei for the selected markers, or by measuring the stained area. The percentage of positive nuclei for selected stainings was determined by first applying automated nuclear segmentation using the cellpose python environment (https://www.cellpose.org/), followed by manual analysis and scoring of the stained nuclei. The positive stained areas were defined and measured via setting common thresholds in Fiji. At least 10 organoids per clone (n=number of clones) and experiment (N=number of experiments) were analysed for quantifications. At least N=3 independent experiments were performed. Intensity measurements of ciliary proteins were performed on unprocessed images (raw data) as described before^18–21, 54^. Analyses of ciliary density were performed by manually counting cilia axonemes over the number of basal bodies (γ-Tubulin). Whenever possible, at least 100 cilia per clone were used for intensity, length and density measurements. After quantifications were performed, representative images were processed by means of background subtraction and contrast settings via Adobe Photoshop CS2.

Data are presented as mean ± SEM or as median with quartiles. Two-tailed t test with Welch’s correction was performed for all data in which two datasets were compared. Two-tailed *t* test Analysis of variance (ANOVA) and Tukey honest significance difference (HSD) tests were used for all data in which more than two datasets were compared. Following statistical significances were considered: \**P* < 0.05, \*\**P* < 0.01, \*\*\**P* < 0.001 and \*\*\*\**P* < 0.0001. All statistical data analysis and graph illustrations were performed using GraphPad Prism (GraphPad Software).

### RNA isolation and qPCR analyses

mESC- and hiPSC-derived organoids were collected at different stages during differentiation and washed with PBS. RNA was isolated using the RNeasy Kit (Qiagen #74104) and RNAse-free DNase Set (Qiagen #79254). Isolated RNA was transcribed into cDNA using the GoScript Reverse Transcriptase Kit (Promega #A5001). For quantitative real-time PCR, 50 ng of cDNA of each sample was used in a Maxima SYBR Green/ROX qPCR Master Mix 2x (Thermo Scientific #K0222). Reactions were run in a Step One Real-Time PCR System Thermal Cycling Block (Applied Biosystems #4376357). Primer pairs are listed in Table S1. The analysis of real-time data was performed using the included StepOne Software version 2.0 and graphs were generated using GraphPad Prism (GraphPad Software).

### Bulk RNAseq analysis on mouse and human spinal organoids

mESC- and hiPSC-derived organoids were collected at different stages during differentiation and washed with PBS. For mouse organoids, pooled organoids from Wt and *Rpgrip1l KO* were collected from 3 independent experiments (N=3). For human organoids, material from two control clones of the PCIi033-A hiPSC line and the UCSFi001-A hiPSC line each (WT: n=4) and from one RPGRIP1L-deficient clone of the PCIi033-A hiPSC line and the UCSFi001-A hiPSC line each (KO: n=2) was collected. RNA was isolated using the RNeasy Kit (Qiagen #74104) and RNAse-free DNase Set (Qiagen #79254). RNA quality was determined by measuring the RNA integrity number (RIN) via the High Sensitive RNA Screen Tape Analysis kit (Agilent Technologies #5067) on the TapeStation system (Agilent Technologies). A RIN above 9.5 was considered as good quality RNA and 250 ng RNA in a total volume of 25 µl was prepared per sample for further procedure. Bulk RNAseq was realized by the Genotyping and Sequencing Core Facility of the Paris Brain Institute (iGenSeq, ICM Paris). RNAseq data processing was performed in Galaxy under supervision of the ARTBio platform (IBPS, Paris).

Paired-end RNA reads were aligned against the *M. musculus mm39* and the *homo sapiens hg38* genome, respectively, by using the STAR aligner (v2.7.10, Galaxy wrapper v4)^128^ and the gene annotations gtf files GRCm39.109 and GRCh38.109, respectively. Quality of sequencing was controlled with fastQC (Galaxy wrapper v0.74+galaxy0) and MultiQC (Galaxy wrapper v1.11+galaxy1)^129^. Gene expression was assessed by counting sequencing reads aligned to gene exons with featureCounts (Galaxy wrapper v2.0.3+galaxy2)^130^. Raw counts were further converted to normalized gene expression values using the log2(tpm+1) transformation where tpm is the count of transcript aligned reads per length of transcript (kb) per million of total mapped reads (Galaxy tool “cpm_tpm_rpk” v0.5.2). Principal Component Analyses (PCA) were performed using r-factominer (v.2.9, Galaxy tool id “high_dimensions_visualisation” v4.3+galaxy0)^131^ and heatmap visualizations were produced with the Galaxy tool “high_dim_heatmap” (v3.1.3+galaxy0) using the normalized gene expression values. Differentially expressed genes (DEGs) were selected from the gene raw read counts using DESeq2 (Galaxy wrapper v2.11.40.8+galaxy0)^132^ and the Benjamini-Hochberg p-adjusted cutoff 0.01. DESeq2 statistical tables were used for generation of Volcano Plots (Galaxy tool id “volcanoplot” v0.0.5)^133^, for goseq tests of overrepresented gene ontology categories (Galaxy wrapper v1.50.0+galaxy0)^134^, as well as for Ensemble of Gene Set Enrichment Analyses (EGSEA) (Galaxy wrapper v1.20.0)^135^.

### Analysis of differential temporal profiles on human spinal organoids

We obtained 62710 gene temporal profiles of expression levels (expressed in log2(tpm+1), each composed of 7 time points in the WT and *RPGRIP1L KO* mutant contexts. We selected genes whose variance of expression across the 14 WT and KO time points is higher than 0.1, and we scaled each gene time point (WT and KO) using the minmax function on the TSPred R package, following which the scaled expression in each series is E’ = (E - E_min_) / (E_max_ - E_min_), the highest expression of series is 1 and the lowest expression of series is 0. Note that since we scaled wild type and mutant expressions together, their relationship is maintained during the minmax scaling.

We next split the data into gene time series of 7 points in WT or KO context, and for each gene, we computed the euclidean distance between the wild type and mutant series. The density function of the euclidean distance was empirically derived from the observed distances. Using this function, we selected genes with the 1 % highest distances between time profiles in wild type or mutant context. Time profiles of selected genes were finally plotted using their unscaled expression values. Since the euclidean distance is not able to capture significant differences between all time series, we repeated the above analysis with 11 other distance metrics, including short time series (sts), dynamic time warping (dtw), time alignment measurement (tam), autocorrelation-based dissimilarity (acf), fourier discrete transform (fourier), compression-based dissimilarity (cdm), complexity-invariant (cid), Pearson’s correlation dissimilarities (cor), integrated periodogram-based dissimilarity (int.per), periodogram-based dissimilarity (per) and Frechet (frechet) distances.

Gene time profiles selected using the various distances metrics are compiled in <u>table</u> [distance mode] which indicates for each gene the metrics that returned a significant distance. In total, <u>we selected 542</u> <u>genes</u> with significant distance between mutant and wild-type time profiles. These profiles are sorted according to the number of metrics returning significant distance (p <0.01). R code used for this analysis is available <u>here</u> for download.

## Acknowledgement

Image acquisition was carried out at the IBPS Imaging Facility. We particularly thank Chloé Chaumeton from the IBPS imaging platform, and Frédéric Eghiaian from Abberior Instruments GmbH, for setting up the STED imaging system. We further thank Jean-François Gilles for his help in setting up the python cellpose environment. We thank the ICV-iPS platform at the Paris Brain Institute (ICM) for the generation and characterization of RPGRIP1L-deficient hiPSC lines, and the iGenSeq sequencing platform at the ICM for reliable sequencing and fast processing of our samples. We are very grateful to Vanessa Ribes and Pascal Gilardi Hebenstreit for providing us with protocols and help in mESCs culture and differentiation approaches. We also thank Jean-François Brunet for kindly providing the Phox2b antibody and the TACGENE team of the Natural History Museum in Paris for providing the Cas9 protein. We thank Estelle Balissat for her help in the preparation of TMEM67 spinal cord organoids and cryosections. We also acknowledge the worldwide contributions of users and developers to the Galaxy project (https://galaxyproject.org/) and all the upstream authors and contributors of the software ecosystem we use and rely on. This work was supported by funding to SSM from the *Fondation pour la Recherche Médicale* (Equipe FRM EQU201903007943), from the *Fondation pour la Recherche sur le Cancer* (ARC; PJA 20171206591), from the *Agence Nationale de la Recherche* (ANR; ERA-NET Neuron project on neurodevelopmental disorders “NDCil”). AW received funding from the *German Research Foundation* (DFG; WI 5451/1-1). ST received funding from the *Agence Nationale de la Recherche* (ANR; ANR-17-CE16-0003-01 and ANR-17-RHUS-0002).

## Author contribution

A.S., S.S.M. and A.W. designed the study, interpreted the data and wrote the manuscript. A.W. & A.S. supervised and conducted the experiments. L. M.-L. & L.B. conducted experiments for the production and analysis of mouse and human spinal organoids. D.C, S.T., A.E. & V.S. designed and conducted experiments to produce iPSC mutant models. C.A. designed and conducted RNAseq analysis. S.N. & M.C. participated in designing the study, interpreting the data and writing the manuscript.

## Ethics statement: Competing interests

We declare that none of the authors have competing financial or non-financial interests.

## Data availability statement

RNA sequencing data will be deposited in an approved, publicly available repository (as requested by Nature Communications) upon acceptance of the manuscript for publication. All other data are available on request.

---

**Supplementary Figure 1:**
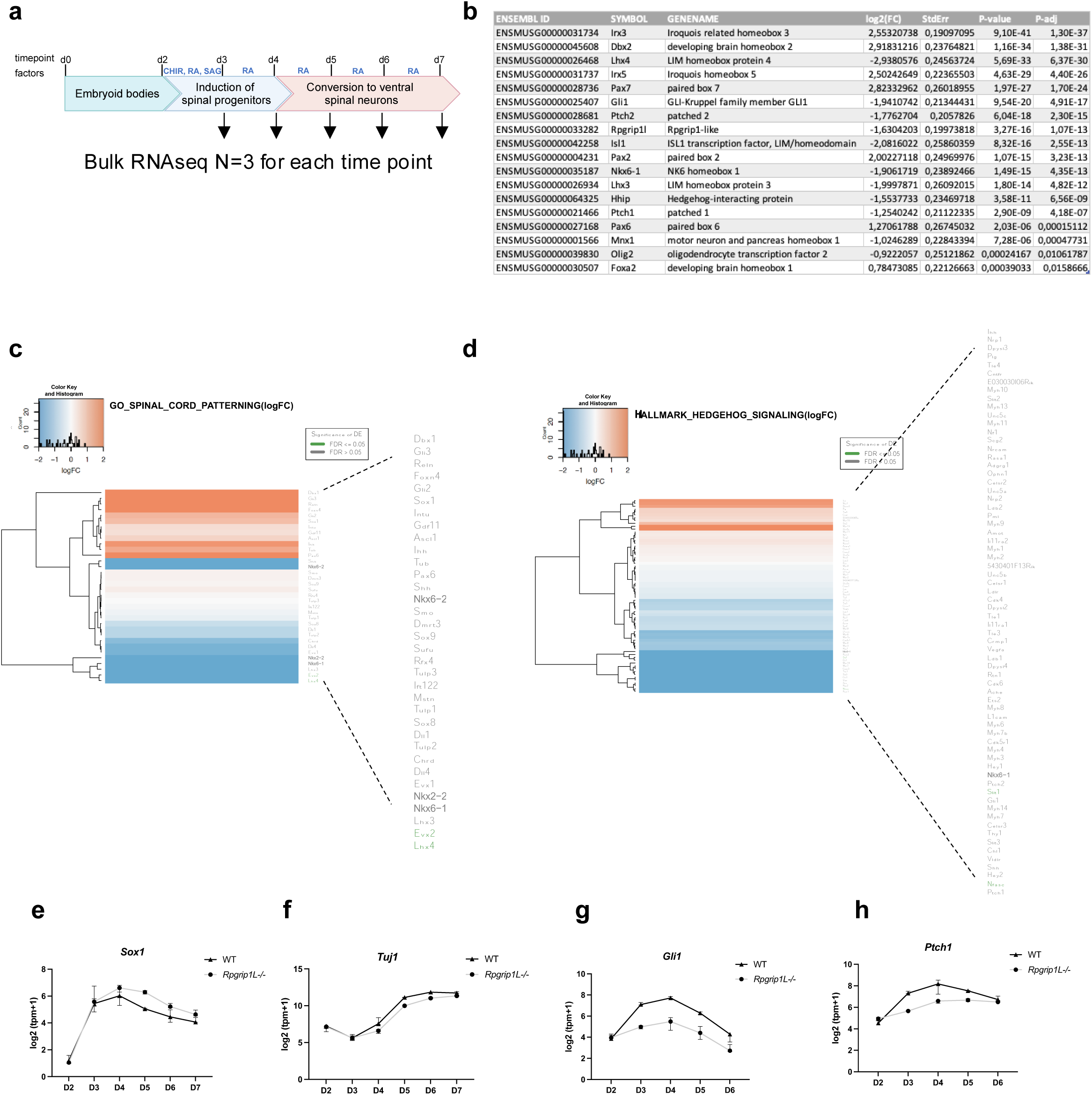
Longitudinal transcriptomic analysis of WT and *Rpgrip1l^-/-^* spinal organoids. **a** Diagram depicting the time points of spinal differentiation chosen for bulk RNAseq analysis. **b** Table listing selected top-differentially expressed genes between WT and *Rpgrip1l^-/-^* day 4 spinal organoids obtained by DEseq2 analysis. **c, d** Ensemble of Gene Set Enrichment Analyses (*EGSEA*) performed on differentially expressed genes between WT and *Rpgrip1l^-/-^* organoids on day 4. Heatmaps ranked 1st for c5 GO Gene Sets (**c**), and h Hallmark Signatures (**d**) are shown. **e-h** Temporal analysis of *Sox1* (NPs), *Tuj1* (early neurons), *Gli1* and *Patched1* (SHH targets) in the course of the differentiation of WT and *Rpgrip1l*^-/-^ organoids. Log2(tpm+1) data from bulk RNASeq analysis are displayed as mean ± SEM (N=3 for each genotype).

**Supplementary Figure 2:**
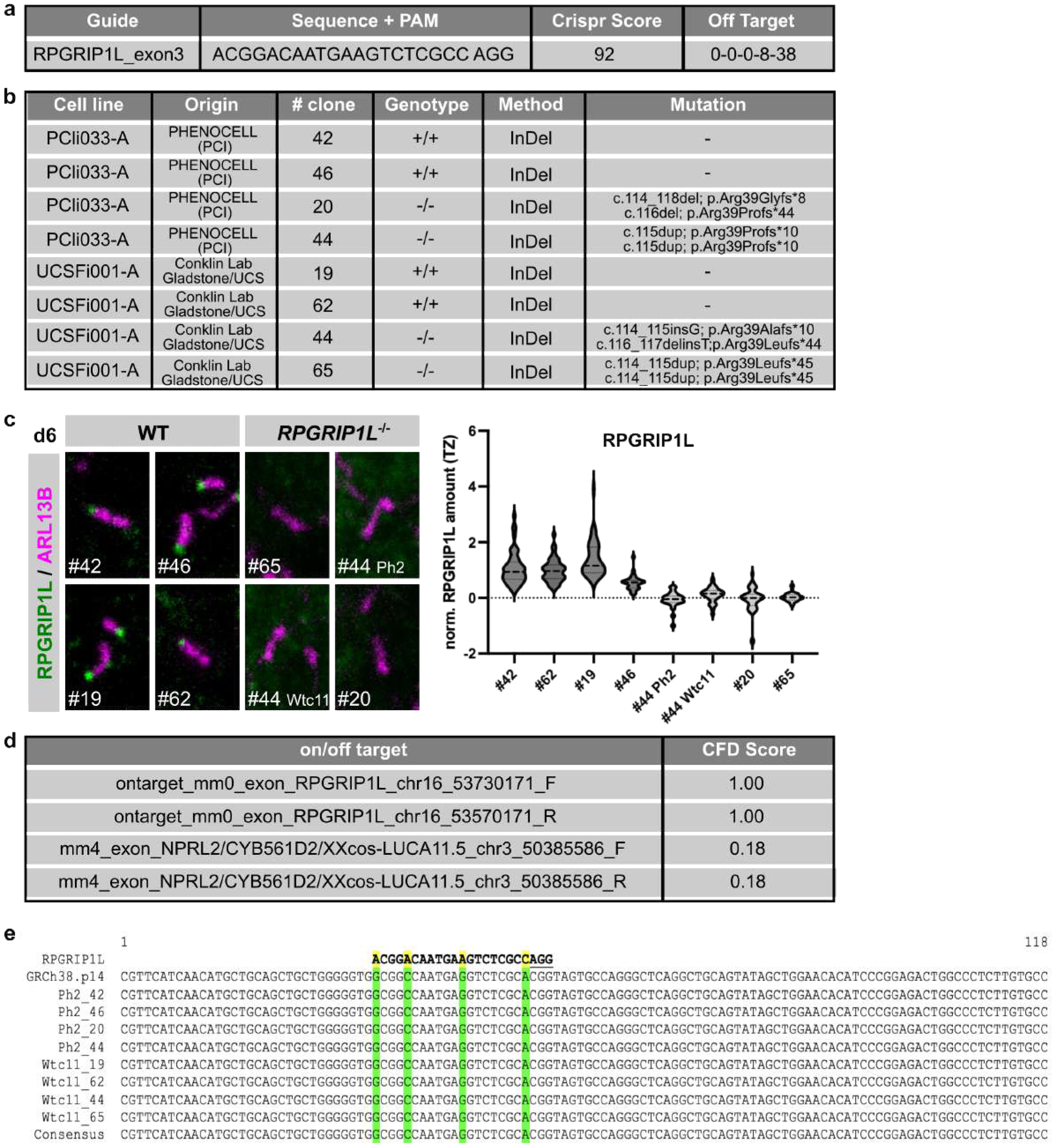
Generation of RPGRIP1L-deficient hiPSC lines. **a** Sequence of the CRISPR-guide that was used for the generation of mutations in exon 3 of *RPGRIP1L*. The Crispr Score corresponds to the CFD specificity score based on the CFD off-target model. The Off Target field indicates the number of off-targets for each number of mismatches (8 off-targets with 3 mismatches and 38 off-targets with 4 mismatches). None of these mismatches are in the 12 bp adjacent to the PAM. **b** List of hiPSC lines that were used in this project. The origin, clone number, genotype and specific mutation are indicated for each clone. **c** Immunofluorescence of RPGRIP1L in WT and RPGRIP1L-deficient cilia in spinal organoids at day 6. Cilia are labeled by ARL13B and RPGRIP1L. Quantifications show the ciliary RPGRIP1L is detectable in WT but not RPGRIP1L-deficient cilia. Data are shown as median with quartiles (number of cilia > 100). **d** Identified off-targets and corresponding CFD scores for the *RPGRIP1L_exon3* guide. **e** Off-target analyses via sequencing of the *NPRL2* locus. Sequence of the CRISPR-guide is indicated above. Bases highlighted in yellow and green indicate mismatches between the guide sequence and the off-target sequence. No mutation in *NPRL2* was detectable around the off-target site in WT and KO hiPSC clones.

**Supplementary Figure 3:**
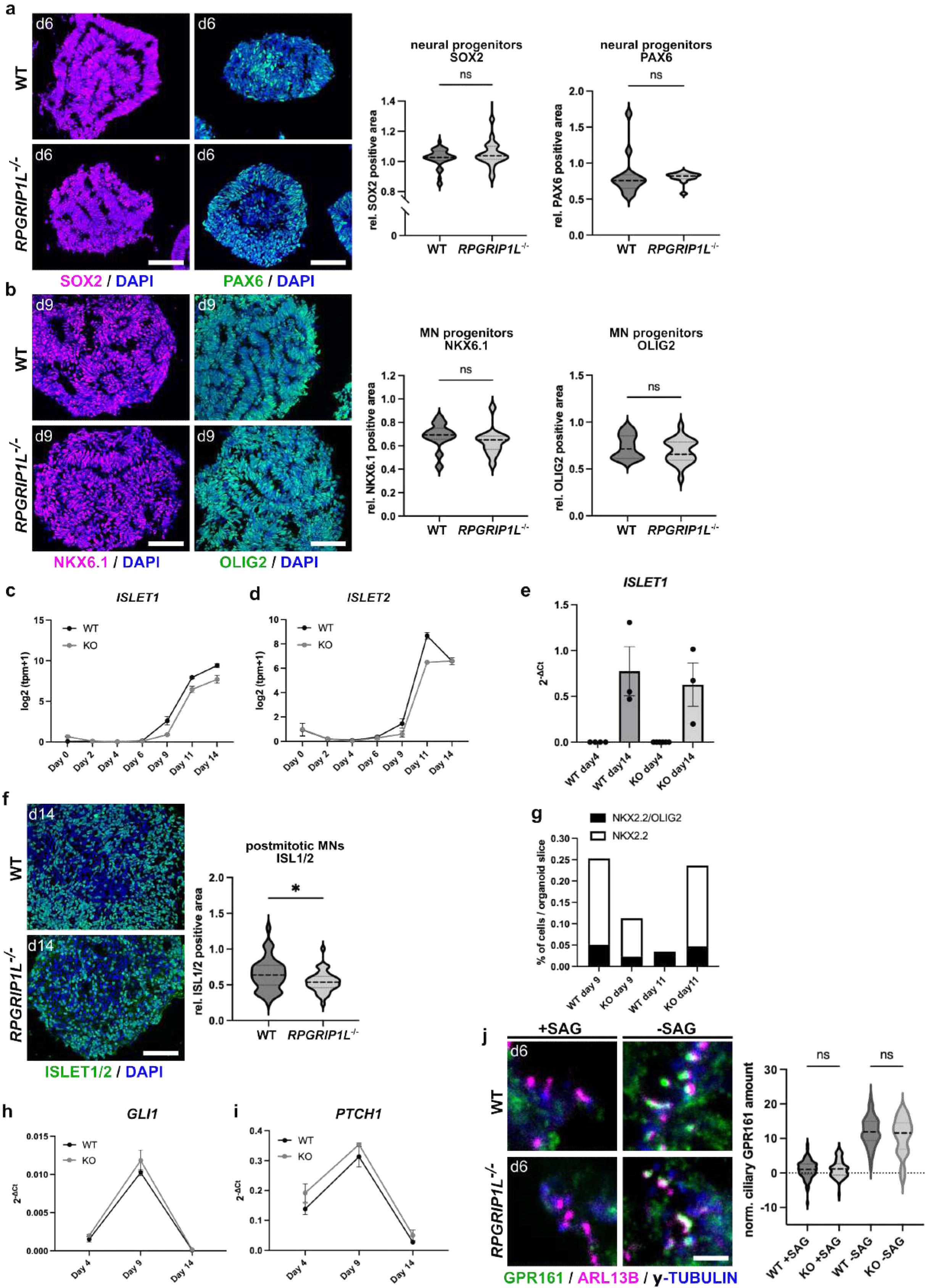
Phenotyping of WT and RPGRIP1L-deficient ventral spinal organoids. **a** Immunofluorescence analysis of neural progenitors in WT and *RPGRIP1L*^-/-^ spinal organoids at day 6. Quantifications show the relative SOX2 and PAX6 positive areas per organoid. Scale bars: 100 µm. Data are shown as median with quartiles (WT: n=4; KO: n=4; N≧3). **b** Immunofluorescence analysis of pMN in WT and *RPGRIP1L*^-/-^ spinal organoids at day 9. Scale bars: 100 µm. Quantifications show relative NKX6.1 and OLIG2 positive areas per organoid. Data are shown as median with quartiles (WT: n=4; KO: n=4; N≧3). **c, d** Log2(tpm+1)-graphs show dynamic expression levels of *ISLET1* (**c**) and *ISLET2* (**d**) in WT and *RPGRIP1L*^-/-^ organoids. Data are shown as mean ± SEM (WT: n=4; KO: n=2). **e** qPCR analysis of *ISLET1* in WT and *RPGRIP1L*^-/-^ organoids at day 4 and day 14 (WT: n=3; KO: n=3). **f** Immunofluorescence analysis of MNs in WT and *RPGRIP1L*^-/-^ spinal organoids at day 14. Scale bar: 100 µm. Quantifications show relative ISLET1/2 positive areas per organoid. Data are shown as median with quartiles. Asterisks denote statistical significance according to unpaired t tests with Welch’s correction (*P < 0.05) (WT: n=4; KO: n=4; N≧3). **g** Quantification of the percentage of NKX2.2-positive and NKX2.2/OLIG2 double-positive cells per organoid section for WT and RPGRIP1L-deficient spinal organoids at day 9 and day 11 (WT: n=4; KO: n=4). Chi-square tests were performed to test for significant differences. **h, i** qPCR analyses of SHH target genes *GLI1* and *PTCH1* in WT and *RPGRIP1L*^-/-^ organoids at different stages. Data are shown as mean ± SEM (WT: n=3; KO: n=3). **j** Immunofluorescence analysis of the ciliary GPR161 amount in WT and *RPGRIP1L*^-/-^ spinal organoids at day 6. Organoids are either treated with SAG to stimulate SHH signaling or DMSO-treated as control. Quantifications show normalized GPR161 amounts within cilia. Scale bar: 2.5 µm. Data are shown as median with quartiles (WT: n=4; KO: n=4).

**Supplementary Figure 4:**
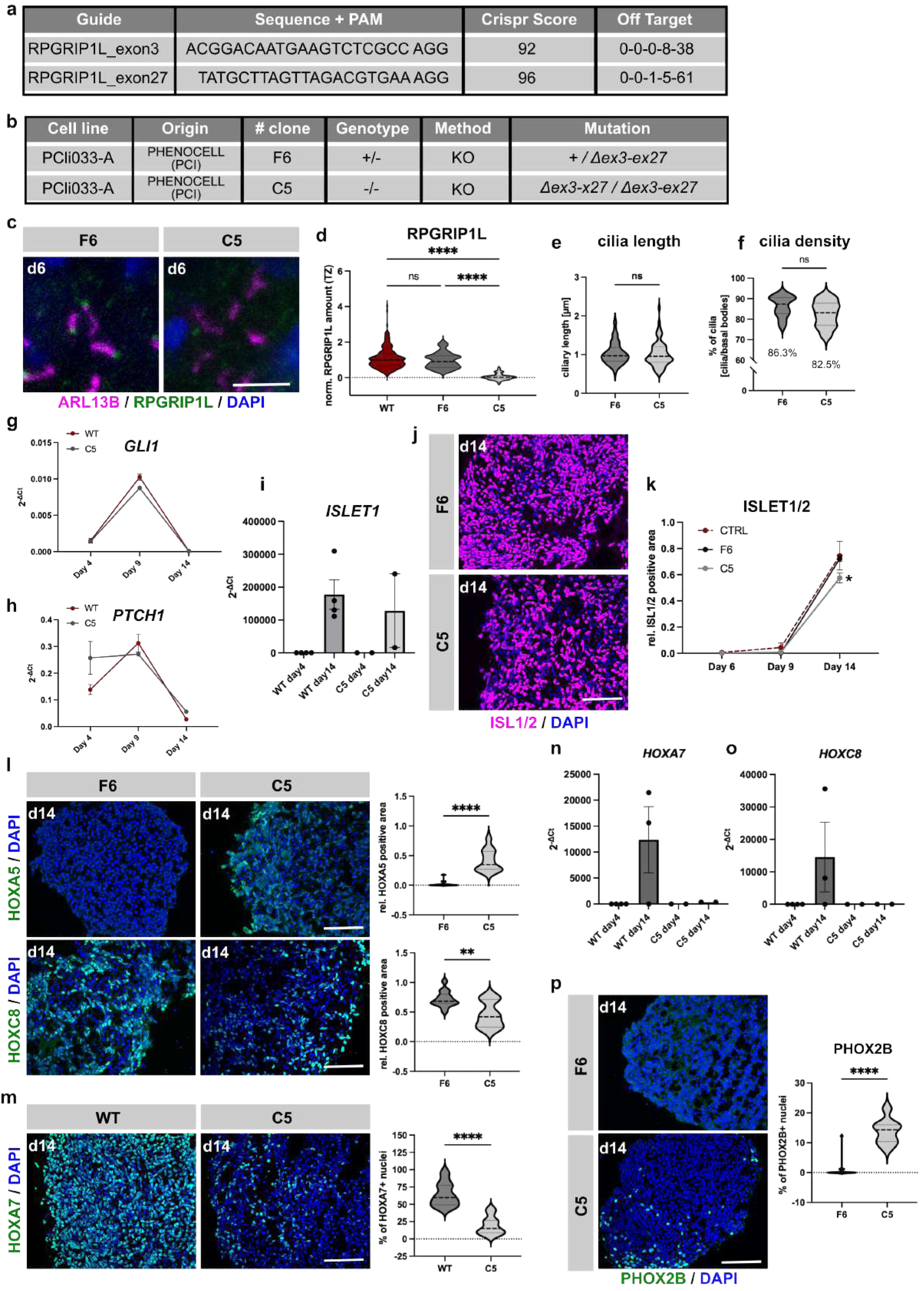
Full-deletion RPGRIP1L hiPSCs show identical phenotypes during differentiation compared to *InDel RPGRIP1L KO* hiPSCs. **a** Sequences of the CRISPR-guides in exon 3 and exon 27 of *RPGRIP1L*, that were used for the generation of a full-deletion hiPSC line. The CRISPR Scores correspond to the CFD specificity scores based on the CFD off-target model. The Off Target field indicates the number of off-targets for each number of mismatches. None of these mismatches are in the 12 bp adjacent to the PAM. **b** hiPSC lines that were used in this project. The origin, clone number, genotype and specific mutation are indicated for both clones. **c** Immunofluorescence of RPGRIP1L in *RPGRIP1L*^+/-^ and *RPGRIP1L*^-/-^ spinal organoids at day 6. Cilia are labeled by ARL13B and RPGRIP1L. Scale bar: 2.5 µm. **d** Quantification of ciliary RPGRIP1L amounts in WT, *RPGRIP1L*^+/-^ and *RPGRIP1L*^-/-^ spinal organoids at day 6. Data are shown as median with quartiles. Asterisks denote statistical significance according to a two-tailed t test Analysis of variance (ANOVA) and Tukey honest significance difference (HSD) (****P < 0.0001) (WT: n=4; F6: N=3; C5: N=3). **e, f** Quantifications show ciliary length and cilia density measurements in *RPGRIP1L*^+/-^ and *RPGRIP1L*^-/-^ organoids at day 6. Data are shown as median with quartiles (F6: N=3; C5: N=3). **g, h** qPCR analyses of SHH target genes *GLI1* and *PTCH1* in WT and *RPGRIP1L*^-/-^ organoids at different time points (WT: n=4; C5: N=3). **i** qPCR analysis of *ISLET1* in WT and *RPGRIP1L*^-/-^ organoids at day 4 and day 14 (WT: n=4; C5: N=2). **j** Immunofluorescence analysis of MNs (ISLET1/2) in *RPGRIP1L*^+/-^ and *RPGRIP1L*^-/-^ spinal organoids. Scale bar: 250µm. **k** Quantifications show the relative ISLET1/2 positive area per WT, *RPGRIP1L*^+/-^ and *RPGRIP1L*^-/-^ spinal organoid over time. Data are shown as mean ± SEM. Asterisks denote statistical significance according to a two-tailed t test Analysis of variance (ANOVA) and Tukey honest significance difference (HSD) (*P < 0.05) (WT: n=4; F6: N=3; C5: N=3). **l** Immunofluorescence analysis of HOXA5 and HOXC8 in *RPGRIP1L*^+/-^ and *RPGRIP1L*^-/-^ spinal organoids at day 14. Quantifications show relative HOXA5 and HOXC8 positive areas per organoid. Data are shown as median with quartiles. Asterisks denote statistical significance according to unpaired t tests with Welch’s correction (**P < 0.001; ****P < 0.0001) (F6: N=3; C5: N=3). **m** Immunofluorescence analysis of HOXA7 in WT and *RPGRIP1L*^-/-^ spinal organoids at day 14. Quantifications show the percentage of HOXA7 positive nuclei per organoid. Data are shown as median with quartiles. Asterisks denote statistical significance according to unpaired t tests with Welch’s correction (****P < 0.0001) (WT: n=4; C5: N=3). **n, o** qPCR analyses of *HOXA7* and *HOXC8* in WT and *RPGRIP1L*^-/-^ organoids at day 4 and day 14 (WT: n=3; C5: N=2). **p** Immunofluorescence analysis of PHOX2B in *RPGRIP1L*^+/-^ and *RPGRIP1L*^-/-^ spinal organoids on day 14. Quantifications show the percentage of PHOX2B positive nuclei per organoid. Data are shown as median with quartiles. Asterisks denote statistical significance according to unpaired t tests with Welch’s correction (****P < 0.0001) (F6: N=3; KO: N=3).

**Supplementary Figure 5:**
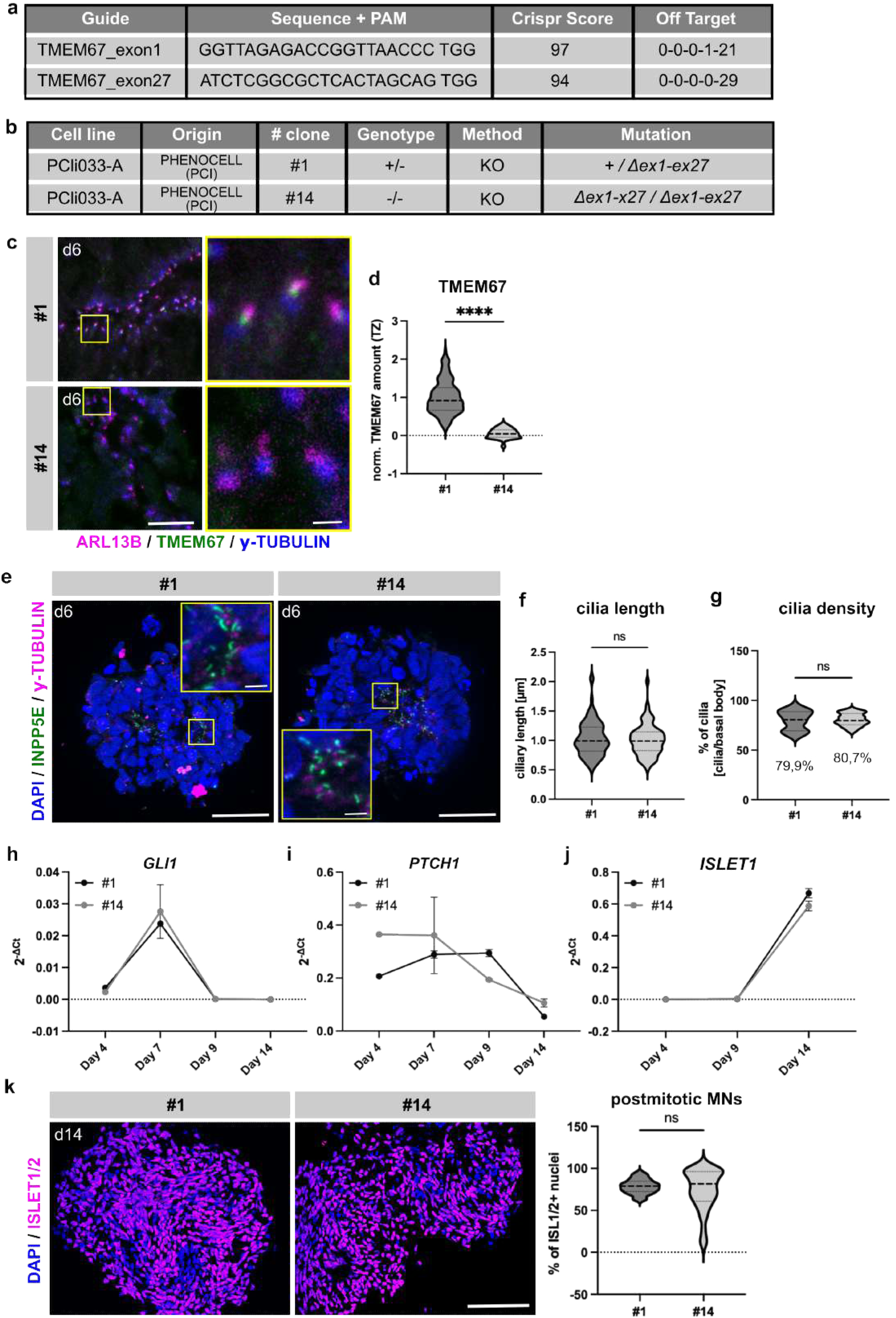
Spinal MN differentiation of TMEM67-deficient hiPSC lines. **a** Sequences of the CRISPR-guides in exon 1 and exon 27, that were used for the generation of a full-deletion hiPSC line. The CRISPR Score corresponds to the CFD specificity score based on the CFD off-target model. The Off Target field indicates the number of off-targets for each number of mismatches. None of these mismatches are in the 12 bp adjacent to the PAM. **b** List of hiPSC lines that were used in this project. The origin, clone number, genotype and specific mutation are indicated for each clone. **c** Immunofluorescence of TMEM67 in *TMEM67*^+/-^ and *TMEM67*^-/-^ spinal organoids at day 6. Cilia are labeled by ARL13B and basal bodies by γ-TUBULIN. TMEM67 is labeled in green. Scale bar: 10 µm. Magnified areas are indicated by yellow rectangles and magnified images are displayed on the lower right. Scale bar: 1 µm. **d** Quantification of the ciliary TMEM67 amount in *TMEM67*^+/-^ and *TMEM67*^-/-^ spinal organoids at day 6. Data are shown as median with quartiles. Asterisks denote statistical significance according to unpaired t tests with Welch’s correction (****P < 0.0001) (#1: N=3; #14: N=3). e Immunofluorescence staining of primary cilia in *TMEM67*^+/-^ and *TMEM67*^-/-^ spinal organoids. Cilia are labeled by INPP5E and basal bodies by γ-TUBULIN. Scale bars: 50 µm. Magnified areas are indicated by yellow rectangles and magnified images are displayed on the lower left. Scale bars: 2 µm. **f, g** Quantification of cilia length and density in *TMEM67*^+/-^ and *TMEM67*^-/-^ spinal organoids at day 6 (#1: N=3; #14: N=3). **h, i, j** qPCR analyses of *GLI1* (**h**), *PTCH1* (**i**) and *ISLET1* (**j**) gene expression profiles in *TMEM67*^+/-^ and *TMEM67*^-/-^ spinal organoids over time. Data are presented as mean ± SEM (#1: N=3; #14: N=3). **k** Immunofluorescence analysis of MNs in *TMEM67*^+/-^ and *TMEM67*^-/-^ spinal organoids at day 14. Scale bar: 150 µm. Quantification shows the percentage of ISLET1/2 positive nuclei per organoid. Data are shown as median with quartiles (#1: N=3; #14: N=3).

**Supplementary Figure 6:**
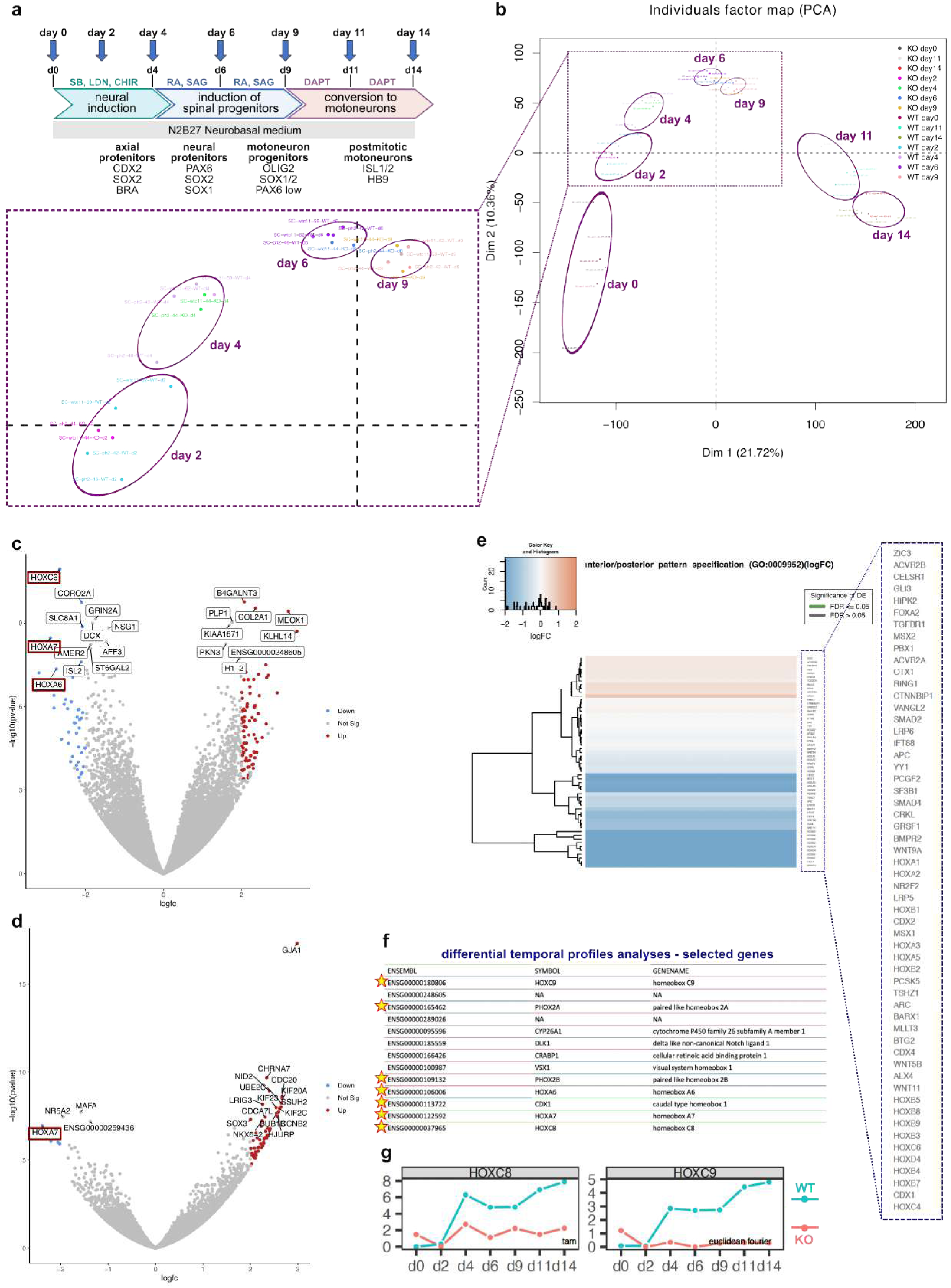
Bulk RNAseq analysis indicates altered antero-posterior patterning in RPGRIP1L-deficient spinal organoids. **a** Schematic summary of the spinal 3D differentiation approach. Samples for bulk RNAseq analyses were collected at indicated time points (WT: n=4; KO: n=2). **b** PCA analyses performed on WT and KO samples show the variation of data over time. The zoom-in on the left shows that WT and KO samples cluster closely together at single time points. **c, d** Volcano plots of RNAseq data on day 11 (**c**) and day 14 (**d**). The threshold for significance was set to 0.01 and the LogFC threshold to 2. **e** EGSEA at day 2 of spinal differentiation shows downregulated *anterior/posterior_pattern_specification* (GO:0009952) in RPGRIP1L-deficient organoids. Genes that correspond to this GO term are depicted on the right. **f** List of genes with different temporal expression profiles based on euclidean distance measurements between WT and *RPGRIP1L KO* organoids over the entire time course of differentiation. Asterisks indicate genes that are related to antero-posterior patterning. **g** Examples of genes with different temporal expression profiles. *HOXC8* was captured via time alignment measurement (tam) and *HOXC9* via euclidean measurement.

**Supplementary Figure 7:**
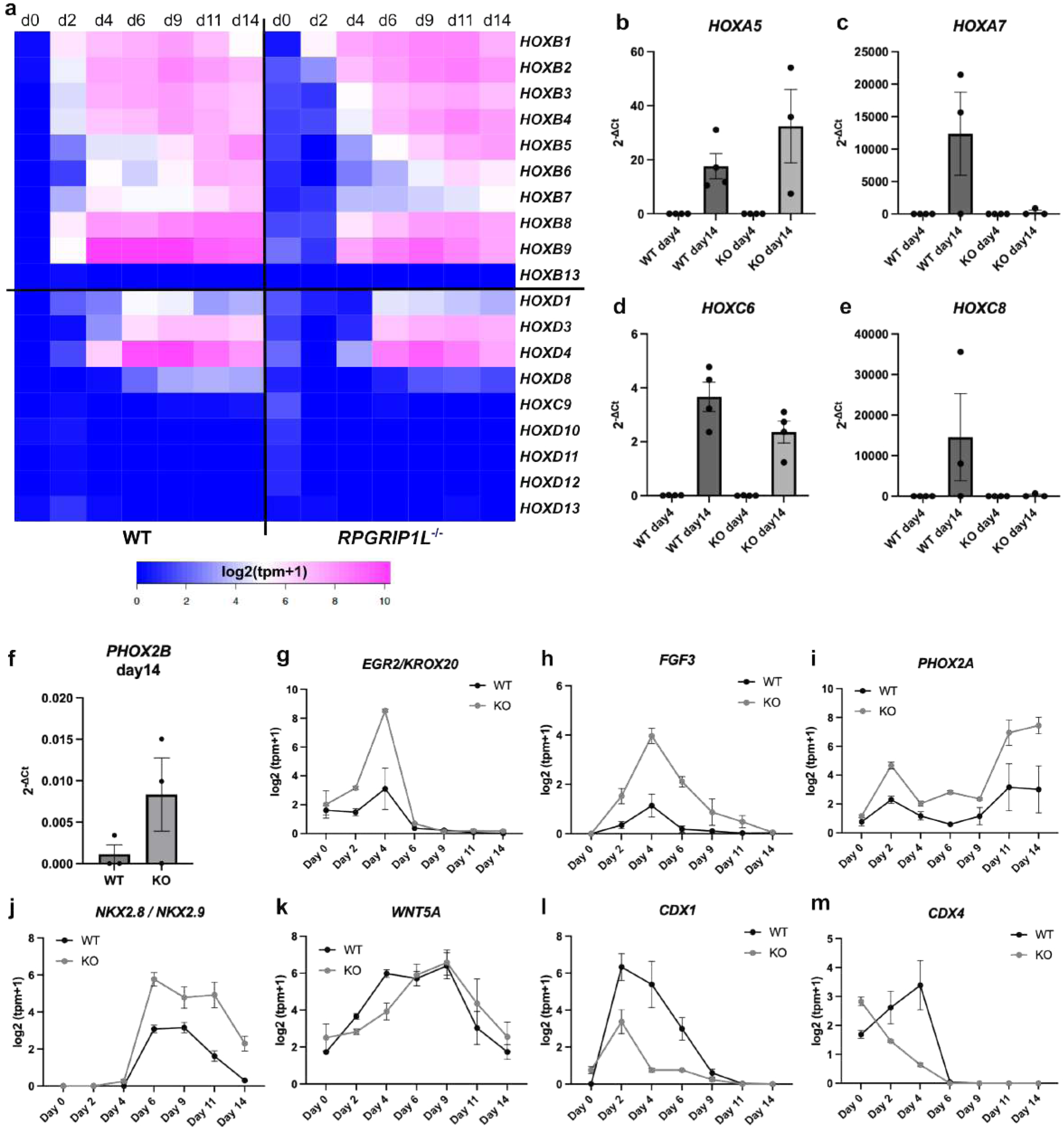
Antero-posterior patterning defects in RPGRIP1L-deficient spinal organoids. **a** Heatmap showing *HOXB* and *HOXD* gene expressions in WT and RPGRIP1L-deficient spinal organoids over time. The graph was generated from log2(tpm+1) files of bulk RNASeq analysis (WT: n=4; KO: n=2). **b-e** qPCR analyses of *HOXA5*, *HOXA7*, *HOXC6* and *HOXC8* in WT and RPGRIP1L-deficient organoids at day 4 and day 14 (WT: n=3; KO: n=3). **f** qPCR analysis of *PHOX2B* in WT and RPGRIP1L-deficient organoids at day 14 (WT: n=3; KO: n=3). **g-m** Log2(tpm+1) graphs show expression profiles of EGR2/KROX20 (**g**) *FGF3* (**h**), *PHOX2A* (**i**), *NKX2.8/NKX2.9* (**j**), *WNT5A* (**k**), *CDX1* (**l**) and *CDX4* (**m**) over time. Data are shown as mean ± SEM (WT: n=4; KO: n=2).

**Supplementary Figure 8:**
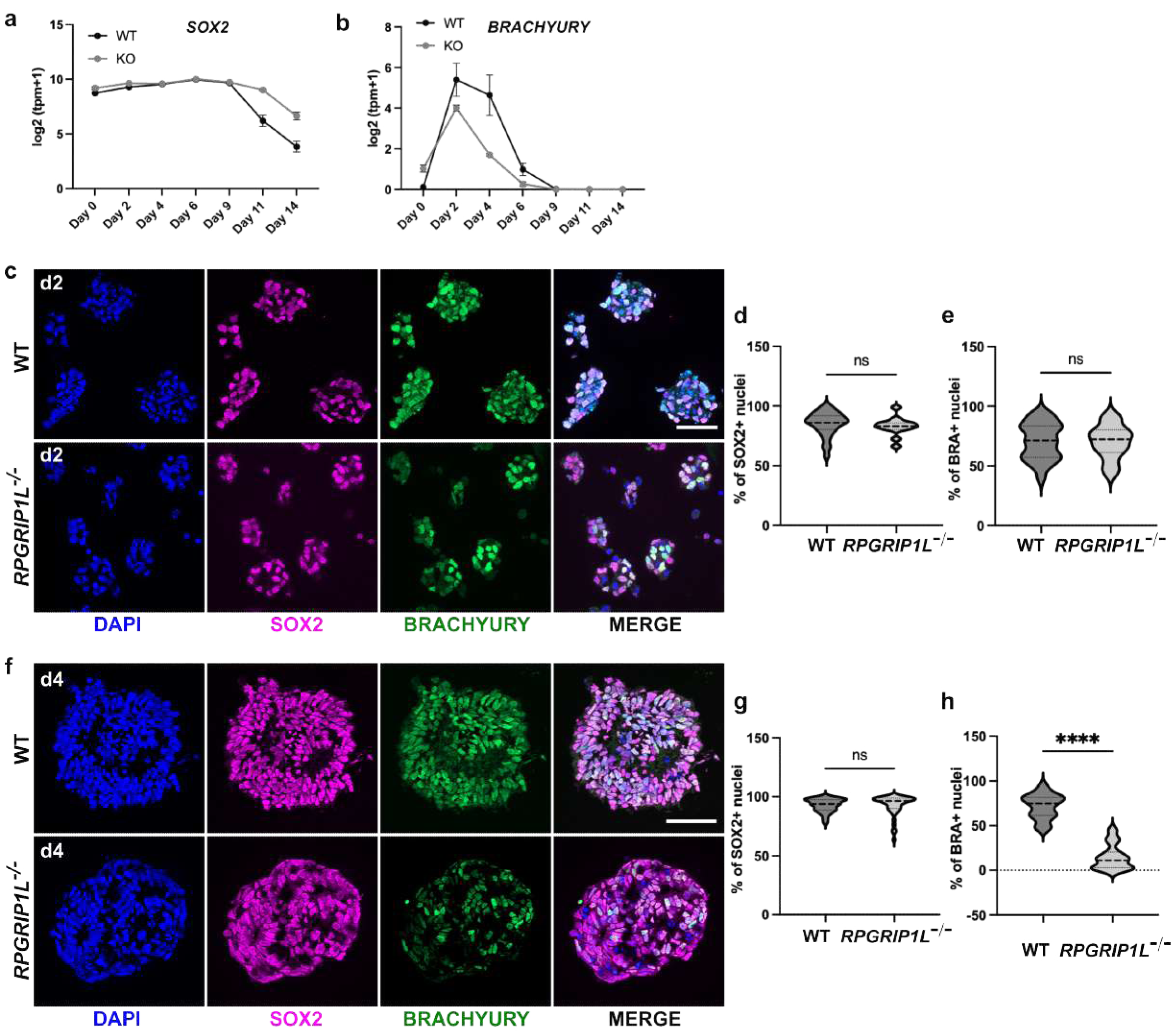
hiPSCs adopt a NMP-like axial fate during early stages of ventral spinal differentiation. **a, b** Log2(tpm+1) graphs generated from bulk RNASeq analyses show the expression profiles of *SOX2* and *BRACHYURY* over time. Data are shown as mean ± SEM (WT: n=4; KO: n=2). **c, f** Immunofluorescence of SOX2 and BRACHYURY in WT and RPGRIP1L-deficient spinal organoids at day 2 (**c**) and day 4 (**f**). Scale bars: 150 µm. **d, e, g, h** Quantifications show the percentage of SOX2 and BRACHYURY positive nuclei per organoid at day 2 and day 4. Data are shown as median with quartiles (WT: n=4; KO: n=4). Asterisks denote statistical significance according to unpaired t tests with Welch’s correction (****P < 0.0001).

**Supplementary Figure 9:**
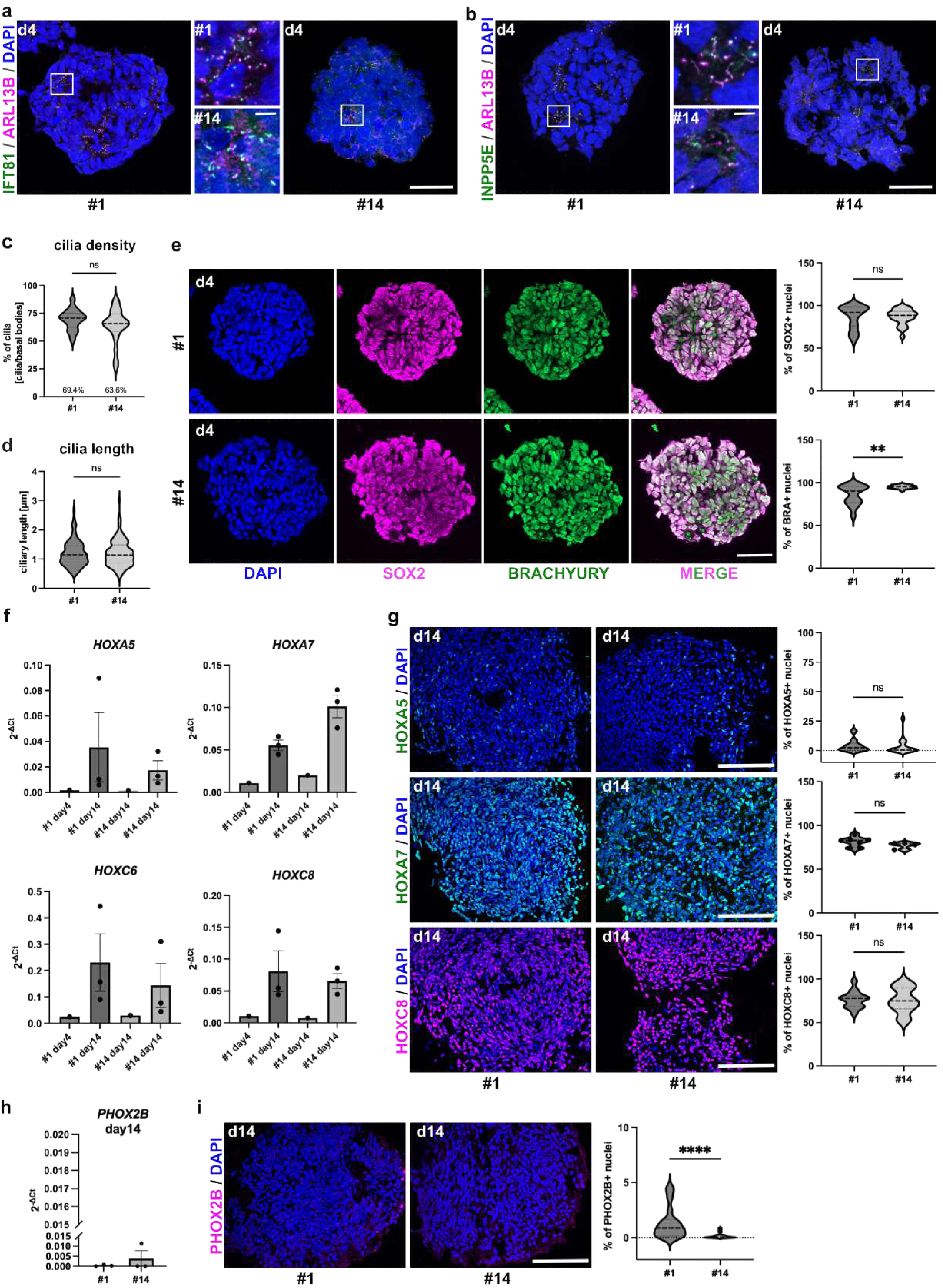
TMEM67-deficient spinal organoids display unaltered axial progenitor specification and correct antero-posterior patterning. **a, b** Immunofluorescence of cilia in *TMEM67*^+/-^ and *TMEM67*^-/-^ spinal organoids at day 4. Cilia are labeled by IFT81and ARL13B (**a**) or by INPP5E and ARL13B (**b**) Scale bars: 50 µm. Magnified areas are indicated by yellow rectangles and magnified images are displayed in the middle. Scale bars: 5 µm. **c, d** Quantification of cilia density and ciliary length in *TMEM67*^+/-^ and *TMEM67*^-/-^ spinal organoids at day 4. Data are shown as median with quartiles (#1: N=3; #14: N=3). **e** Immunofluorescence of SOX2 and BRACHYURY in *TMEM67*^+/-^ and *TMEM67*^-/-^ spinal organoids at day 4. Data are shown as median with quartiles (#1: N=3; #14: N=3). **f** qPCR analyses of *HOXA5*, *HOXA7*, *HOXC6* and *HOXC8* in *TMEM67*^+/-^ and *TMEM67*^-/-^ organoids at day 4 and day 14 (day4: #1 N=1, #14 N=1; day14: #1 N=3, #14: N=3). **g** Immunofluorescence of HOXA5, HOXA7 and HOXC8 in *TMEM67*^+/-^ and *TMEM67*^-/-^ spinal organoids at day 14. Scale bars: 150 µm. Quantifications show the percentages of HOXA5, HOXA7 and HOXC8 positive nuclei per organoid. Data are shown as median with quartiles (#1: N=3; #14: N=3). **h** qPCR analysis of *PHOX2B* in *TMEM67*^+/-^ and *TMEM67*^-/-^ spinal organoids at day 14 (#1: N=3; #14: N=3). **i** Immunofluorescence of PHOX2B in *TMEM67*^+/-^ and *TMEM67*^-/-^ spinal organoids at day 14. Scale bar: 150 µm. Quantifications show the percentage ofPHOX2B positive nuclei per organoid. Data are shown as median with quartiles (#1: N=3; #14: N=3). Asterisks denote statistical significance according to unpaired t tests with Welch’s correction (****P < 0.0001).

**Supplementary Figure 10:**
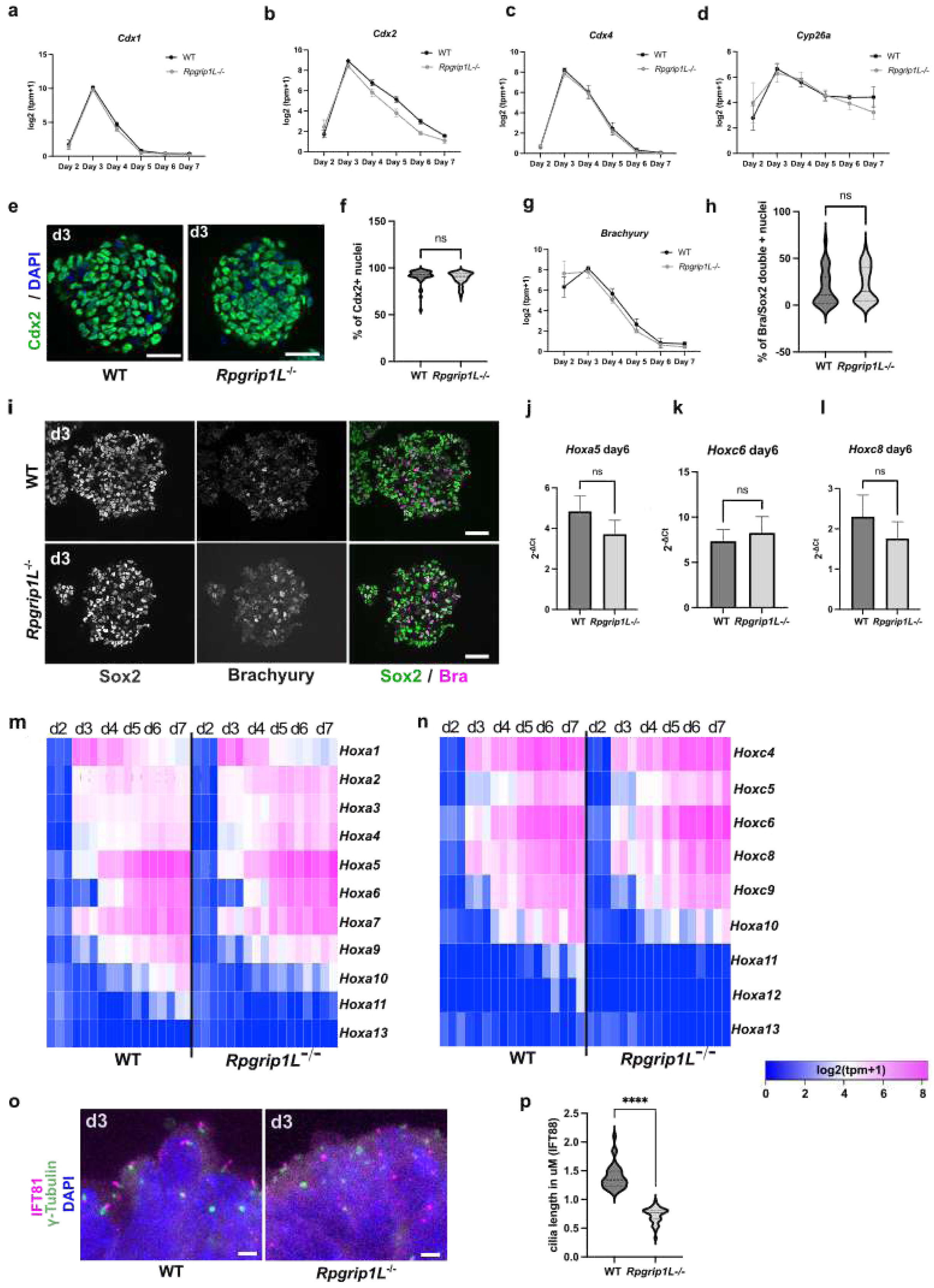
Mouse Rpgrip1l-deficient spinal organoids do not display spinal to cranial specification defects. **a-d, g** Temporal analysis of *Cdx1, Cdx2, Cdx4* (spinal identity) *Cyp26a1* (cranial identity) and *Brachyury* (NMPs) expression in the course of the differentiation of WT and *Rpgrip1l*^-/-^ organoids. Log2(tpm+1) data from bulk RNASeq analysis are displayed as mean ± SEM (N=3 for each genotype). **e** Immunofluorescence for Cdx2 on sections from WT and *Rpgrip1l*^-/-^ organoids at day 3. Scale bar, 10 μm. **f** Percentage of nuclei positive for Cdx2 in WT and *Rpgrip1l*^-/-^ day 3 organoids. Data are shown as violin plots, median quartiles are indicated as dotted lines. n organoids analyzed ≧6, N=3 independent experiments for each genotype. **h** Percentage of nuclei double positive for Sox2 and Brachyury in WT and *Rpgrip1l*^-/-^ day 3 organoids. Data are shown as violin plots, median quartiles are indicated as dotted lines. n organoids analyzed ≧6, N=3 independent experiments for each genotype. **i** Immunofluorescence of Sox2 and Brachyury in WT and *Rpgrip1l*^-/-^ day 3 organoids. Scale bar, 10 μm. **j-l** qPCR analyses of indicated genes from WT and *Rpgrip1l*^-/-^ day 6 organoids. Data are displayed as mean ± SEM (N=4 independent experiments for each genotype). **m, n** Heatmap depicting the expression log2(tpm+1) of selected *Hox* genes in WT and *Rpgrip1l KO* spinal organoids over time. N=3 experimental replicates for each genotype. **o** Immunofluorescence for the indicated ciliary markers on sections of day 3 WT and *Rpgrip1l*^-/-^ organoids**. p** Quantifications of ciliary stainings. Data shown are mean ± SEM. n≧20 cilia from n≧3 organoids were analyzed per genotype. N≧2 Number of independent experiments ***P < 0.001, ****P < 0.0001 (unpaired t tests with Welch’s correction). Scale bar, 2 μm.

**Supplementary Table S1.**
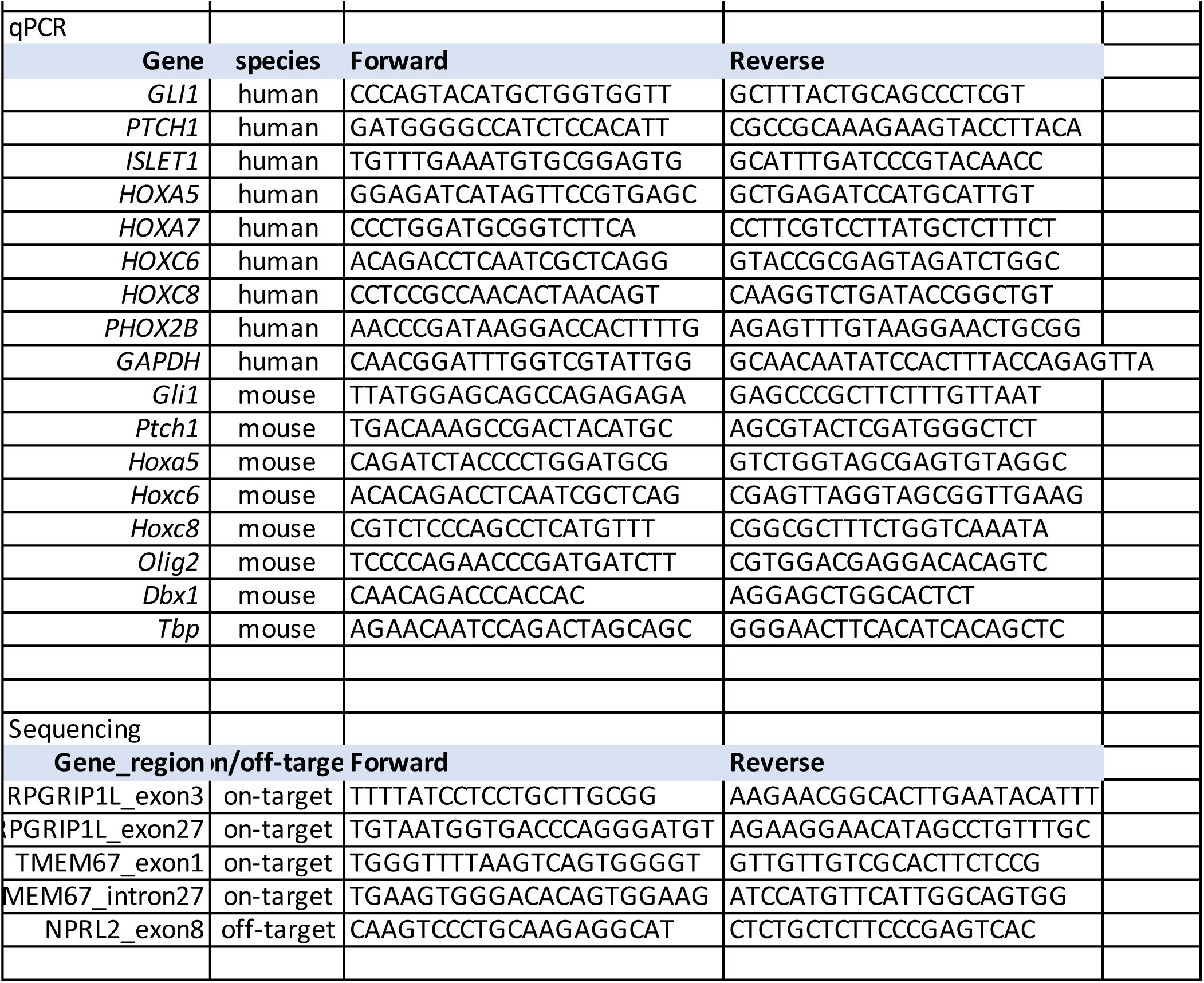

**Supplementary Table 2.**

| primary Antibody | host | company | reference |
| --- | --- | --- | --- |
| Ac3 | rabbit | Abcam | ab125093 |
| Arl13b | mouse IgG2a | Antibody Incorporated | 75-287 |
| Brachyury | rabbit | Cell Signaling/Ozyme | 81694 |
| Brachyury | goat | R&D systems | AF2085 |
| Cdx2 | rabbit | Abcam | ab76541 |
| Gpr161 | rabbit | Proteintech | 13398-1-AP |
| Hoxa5 | rabbit | Sigma | HPA029319 |
| Hoxa7-CoraLite® | mouse IgG1 | Proteintech | CL488-67112 |
| Hoxc8 | rabbit | Sigma | HPA028911 |
| Ift81 | rabbit | Proteintech | 11744-1-AP |
| Ift88 | rabbit | Proteintech | 13967-1-AP |
| Inpp5e | rabbit | Proteintech | 17797-1-AP |
| Isl1/2 | mouse IgG2b | DSHB | 39.4D5 |
| Nkx2.2 | mouse IgG2b | DSHB | 74.5A5 |
| Nkx6.1 | mouse IgG1 | DSHB | F55A12 |
| Olig2 | rabbit | Millipore | AB9610 |
| Pax6 | rabbit | BioLegend | PRB-278P |
| Pax6 | mouse IgG1 | DSHB | PAX6 |
| FoxA2 | mouse IgG1 | DSHB | 4C7 |
| Phox2b | rabbit | kindly provided by JF Brunet | / |
| RPGRIP1L | rabbit | homemade anti-human | against human RID domain (Uniprot A0A087WX34) |
| Rpgrip1l | rabbit | homemade anti-mouse | against mouse RID-domain (PMID: 26150391) |
| Sox2 | rabbit | Millipore | AB5603 |
| Sox2 | mouse | Abcam | ab79351 |
| Tmem67 | rabbit | Proteintech | 13975-1-AP |
| Tuj1 | mouse IgG2a | BioLegend | MMS-435P |
| γ-Tubulin | mouse IgG1 | Sigma | T6557 |
| secondary Antibody | host | conjugate | company |
| Mouse IgG1 | goat | Alexa488 | Molecular Probes |
| Mouse IgG1 | goat | Alexa633 | Molecular Probes |
| Mouse IgG2a | goat | Alexa488 | Molecular Probes |
| Mouse IgG2a | goat | Alexa594 | Molecular Probes |
| Mouse IgG2a | goat | Alexa633 | Molecular Probes |
| Mouse IgG2b | goat | Alexa488 | Molecular Probes |
| Mouse IgG2b | goat | Alexa594 | Molecular Probes |
| Mouse IgG2b | goat | Alexa633 | Molecular Probes |
| Rabbit | goat | Alexa488 | Molecular Probes |
| Rabbit | goat | Alexa594 | Molecular Probes |
| Rabbit | goat | Alexa633 | Molecular Probes |

## References

1. Bachmann-Gagescu, R. et al. Healthcare recommendations for Joubert syndrome. Am J Med Genet A. 182, 229–249. (2020).

2. Reiter, J. & Leroux, M. Genes and molecular pathways underpinning ciliopathies. Nat. Rev. Mol. Cell Biol. 18, 533–547. (2017).

3. Brancati, F., Dallapiccola, B. & Valente, E. Joubert Syndrome and related disorders. Orphanet J Rare Dis. 5, 20. (2010).

4. Goetz, S. & Anderson, K. The primary cilium: a signalling centre during vertebrate development. Nat. Rev. Genet. 11, 331–344. (2010).

5. Spassky, N. et al. Primary cilia are required for cerebellar development and Shh-dependent expansion of progenitor pool. Dev Biol. 317, 246–259. (2008).

6. Andreu-Cervera, A., Catala, M. & Schneider-Maunoury, S. Cilia, ciliopathies and hedgehog-related forebrain developmental disorders. Neurobiol. Dis. 150, 105236 (2021).

7. Andreu-Cervera, A. et al. The ciliopathy gene ftm/rpgrip1l controls mouse forebrain patterning via region-specific modulation of hedgehog/gli signaling. J. Neurosci. 39, 2398–2415 (2019).

8. Abdelhamed, Z. et al. The ciliary Frizzled-like receptor Tmem67 regulates canonical Wnt/β-catenin signalling in the developing cerebellum via Hoxb5. Sci Rep. 9, 5446. (2019).

9. Lancaster, M. et al. Defective Wnt-dependent cerebellar midline fusion in a mouse model of Joubert syndrome. Nat. Med. 17, 726–731. (2011).

10. Basten, S. & Giles, R. Functional aspects of primary cilia in signaling, cell cycle and tumorigenesis. Cilia 2, 6. (2013).

11. Delous, M. et al. The ciliary gene RPGRIP1L is mutated in cerebello-oculo-renal syndrome (Joubert syndrome type B) and Meckel syndrome. Nat. Genet. 39, 875–881. (2007).

12. Arts, H. et al. Mutations in the gene encoding the basal body protein RPGRIP1L, a nephrocystin-4 interactor, cause Joubert syndrome. Nat. Genet. 39, 882–888. (2007).

13. Doherty, D. et al. Mutations in 3 genes (MKS3, CC2D2A and RPGRIP1L) cause COACH syndrome (Joubert syndrome with congenital hepatic fibrosis). J. Med. Genet. 47, 8–21. (2010).

14. Wolf, M. et al. Mutational analysis of the RPGRIP1L gene in patients with Joubert syndrome and nephronophthisis. Kidney Int. 72, 1520–1526. (2007).

15. Vierkotten, J., Dildrop, R., Peters, T., Wang, B. & Rüther, U. Ftm is a novel basal body protein of cilia involved in Shh signalling. Development 134, 2569–2577. (2007).

16. Besse, L. et al. Primary cilia control telencephalic patterning and morphogenesis via Gli3 proteolytic processing. Development 138, 2079–2088. (2011).

17. Shi, X. et al. Super-resolution microscopy reveals that disruption of ciliary transition-zone architecture causes Joubert syndrome. Nat. Cell Biol. 19, 1178–1188. (2017).

18. Wiegering, A. et al. Cell type-specific regulation of ciliary transition zone assembly in vertebrates. EMBO J. 37, e97791. (2018).

19. Wiegering, A. et al. Rpgrip1l controls ciliary gating by ensuring the proper amount of Cep290 at the vertebrate transition zone. Mol. Biol. Cell 32, 675–689. (2021).

20. Gerhardt, C. et al. The transition zone protein Rpgrip1l regulates proteasomal activity at the primary cilium. J. Cell Biol. 210, 115–133. (2015).

21. Struchtrup, A., Wiegering, A., Stork, B., Rüther, U. & Gerhardt, C. The ciliary protein RPGRIP1L governs autophagy independently of its proteasome-regulating function at the ciliary base in mouse embryonic fibroblasts. Autophagy 14, 567–583. (2018).

22. Mahuzier, A. et al. Dishevelled stabilization by the ciliopathy protein Rpgrip1l is essential for planar cell polarity. J. Cell Biol. 198, 927–940. (2012).

23. Gerhardt, C., Wiegering, A., Leu, T. & Rüther, U. Control of Hedgehog signalling by the cilia-regulated proteasome. J. Dev. Biol. 4, 27. (2016).

24. McDonald, A. & Wijnholds, J. Retinal Ciliopathies and Potential Gene Therapies: A Focus on Human iPSC-Derived Organoid Models. International Journal of Molecular Sciences 25, 2887. (2024).

25. Schembs, L. et al. The ciliary gene INPP5E confers dorsal telencephalic identity to human cortical organoids by negatively regulating Sonic hedgehog signaling. Cell Reports 39, 110811. (2022).

26. Boutaud, L. et al. 2D and 3D Human Induced Pluripotent Stem Cell-Based Models to Dissect Primary Cilium Involvement during Neocortical Development. J Vis Exp. 181 (2022).

27. Cruz, N. et al. Modelling ciliopathy phenotypes in human tissues derived from pluripotent stem cells with genetically ablated cilia. Nat Biomed Eng. 6, 463–475. (2022).

28. Willaredt, M. et al. A crucial role for primary cilia in cortical morphogenesis. J Neurosci. 28, 12887–12900. (2008).

29. Juric-Sekhar, G., Adkins, J., Doherty, D. & Hevner, R. Joubert syndrome: brain and spinal cord malformations in genotyped cases and implications for neurodevelopmental functions of primary cilia. Acta Neuropathol. 123, 695–709. (2012).

30. ten Donkelaar, H., Hoevenaars, F. & Wesseling, P. A case of Joubert’s syndrome with extensive cerebral malformations. Clin Neuropathol. 19, 85–93. (2000).

31. Abdelhamed, Z. et al. The Meckel-Gruber syndrome protein TMEM67 controls basal body positioning and epithelial branching morphogenesis in mice via the non-canonical Wnt pathway. Dis Model Mech. 8, 527–541. (2015).

32. Abdelhamed, Z. et al. Variable expressivity of ciliopathy neurological phenotypes that encompass Meckel-Gruber syndrome and Joubert syndrome is caused by complex de-regulated ciliogenesis, Shh and Wnt signalling defects. Hum Mol Genet. 22, 1358–1372. (2013).

33. Adams, M. et al. A meckelin–filamin A interaction mediates ciliogenesis. Hum. Mol. Genet. 21, 1272–1286. (2012).

34. Czarnecki, P.G. & Shah, J.V. The ciliary transition zone: from morphology and molecules to medicine. Trends in Cell Biol. 22, 201–210. (2012).

35. Bangs, F. & Anderson, K. Primary Cilia and Mammalian Hedgehog Signaling. Cold Spring Harb. Perspect. Biol. 9, pii: a028175. (2017).

36. Briscoe, J. & Ericson, J. Specification of neuronal fates in the ventral neural tube. *J*. Curr Opin Neurobiol. 11, 43–49. (2001).

37. Balaskas, N. et al. Gene regulatory logic for reading the Sonic Hedgehog signaling gradient in the vertebrate neural tube. Cell 148, 273–284. (2012).

38. Ribes, V. et al. Distinct Sonic Hedgehog signaling dynamics specify floor plate and ventral neuronal progenitors in the vertebrate neural tube. Genes Dev. 24, 1186–1200. (2010).

39. Dessaud, E. et al. Dynamic assignment and maintenance of positional identity in the ventral neural tube by the morphogen sonic hedgehog. PLoS Biol. 8, e1000382. (2010).

40. Dessaud, E. et al. Interpretation of the sonic hedgehog morphogen gradient by a temporal adaptation mechanism. Nature 450, 717–720. (2007).

41. Lek, M. et al. A homeodomain feedback circuit underlies step-function interpretation of a Shh morphogen gradient during ventral neural patterning. Development 137, 4051–4060. (2010).

42. Huangfu, D. et al. Hedgehog signalling in the mouse requires intraflagellar transport proteins. Nature 426, 83–87. (2003).

43. Haycraft, C. et al. Gli2 and Gli3 localize to cilia and require the intraflagellar transport protein polaris for processing and function. PLoS Genet. 1, e53 (2005).

44. Wichterle, H., Lieberam, I., Porter, J. & Jessell, T. Directed differentiation of embryonic stem cells into motor neurons. Cell 110, 385–397. (2002).

45. Duval, N. et al. BMP4 patterns Smad activity and generates stereotyped cell fate organization in spinal organoids. Development 146, dev175430. (2019).

46. Holzner, M., Wutz, A. & Di Minin, G. Applying Spinal Cord Organoids as a quantitative approach to study the mammalian Hedgehog pathway. PLoS One 19, e0301670. (2024).

47. Maury, Y. et al. Combinatorial analysis of developmental cues efficiently converts human pluripotent stem cells into multiple neuronal subtypes. Nature Biotechnology 33, 89–96. (2015).

48. Mouilleau, V. et al. Dynamic extrinsic pacing of the HOX clock in human axial progenitors controls motor neuron subtype specification. Development 146, dev194514. (2021).

49. Ericson, J., Briscoe, J., Rashbass, P., van Heyningen, V. & Jessell, T. Graded sonic hedgehog signaling and the specification of cell fate in the ventral neural tube. Cold Spring Harb. Symp. Quant. Biol. 62, 451–466. (1997).

50. Pattyn, A. et al. Coordinated temporal and spatial control of motor neuron and serotonergic neuron generation from a common pool of CNS progenitors. Genes Dev. 17, 729–737. (2003).

51. 51. Jang, S., Gunmit, E. & Wichterle, H. Human-specific progenitor sub-domain contributes to extended neurogenesis and increased motor neuron production. bioRxiv (2022).

52. Rayon, T., Maizels, R., Barrington, C. & Briscoe, J. Single-cell transcriptome profiling of the human developing spinal cord reveals a conserved genetic programme with human-specific features. Development 148, dev199711. (2021).

53. Briscoe, J. et al. Homeobox gene Nkx2.2 and specification of neuronal identity by graded Sonic hedgehog signalling. Nature 398, 622–627. (1999).

54. Garcia-Gonzalo, F. et al. A transition zone complex regulates mammalian ciliogenesis and ciliary membrane composition. Nat. Genet. 43, 776–784. (2011).

55. Williams, C. et al. MKS and NPHP modules cooperate to establish basal body/transition zone membrane associations and ciliary gate function during ciliogenesis. J. Cell Biol. 192, 1023–1041. (2011).

56. Jensen, V. et al. Formation of the transition zone by Mks5/Rpgrip1L establishes a ciliary zone of exclusion (CIZE) that compartmentalises ciliary signalling proteins and controls PIP2 ciliary abundance. EMBO J. 34, 2537–2556. (2015).

57. Garcia-Gonzalo, F. et al. Phosphoinositides Regulate Ciliary Protein Trafficking to Modulate Hedgehog Signaling. Dev. Cell 34, 400–409. (2015).

58. Mukhopadhyay, S. et al. The ciliary G-protein-coupled receptor Gpr161 negatively regulates the Sonic hedgehog pathway via cAMP signaling. Cell 152, 210–223. (2013).

59. Pal, K. et al. Smoothened determines β-arrestin-mediated removal of the G protein-coupled receptor Gpr161 from the primary cilium. J Cell Biol. 212, 861–875. (2016).

60. Cambray, N. & Wilson, V. Two distinct sources for a population of maturing axial progenitors. Development. Development. 134, 2829–2840. (2007).

61. Cambray, N. & Wilson, V. Axial progenitors with extensive potency are localised to the mouse chordoneural hinge. Development 129, 4855–4866. (2002).

62. Dasen, J., Liu, J. & Jessell, T. Motor neuron columnar fate imposed by sequential phases of Hox-c activity. Nature 425, 926–933. (2003).

63. Deschamps, J. & Duboule, D. Embryonic timing, axial stem cells, chromatin dynamics, and the Hox clock. Genes Dev. 31, 1406–1416. (2017).

64. Miller, A. & Dasen, J. Establishing and maintaining Hox profiles during spinal cord development. Semin Cell Dev Biol. 152-153, 44–57. (2024).

65. Graham, A., Maden, M. & Krumlauf, R. The murine Hox-2 genes display dynamic dorsoventral patterns of expression during central nervous system development. Development 112, 255–264. (1991).

66. Pattyn, A., Hirsch, M., Goridis, C. & Brunet, J. Control of hindbrain motor neuron differentiation by the homeobox gene Phox2b. Development 127, 1349–1358. (2000).

67. Ahn, D., Ruvinsky, I., Oates, A., Silver, L. & Ho, R. tbx20, a new vertebrate T-box gene expressed in the cranial motor neurons and developing cardiovascular structures in zebrafish. Mech Dev. 95, 253–258. (2000).

68. Guthrie, S. Patterning and axon guidance of cranial motor neurons. Nat Rev Neurosci. 8, 859–871. (2007).

69. Seitanidou, T., Schneider-Maunoury, S., Desmarquet, C., Wilkinson, D. & Charnay, P. Krox-20 is a key regulator of rhombomere-specific gene expression in the developing hindbrain. Mechanisms of Development 65, 31–42 (1997).

70. Schneider-Maunoury, S., Gilardi-Hebenstreit, P. & Charnay, P. How to build a vertebrate hindbrain. Lessons from genetics. C R Acad Sci III. 321, 819–834. (1998).

71. Frank, D. & Sela-Donenfeld, D. Hindbrain induction and patterning during early vertebrate development. Cell Mol Life Sci. 76, 941–960. (2019).

72. Aragon, F. & Pujades, C. FGF signaling controls caudal hindbrain specification through Ras-ERK1/2 pathway. BMC Dev Biol. 9 (2009).

73. Walshe, J., Maroon, H., McGonnell, I., Dickson, C. & Mason, I. Establishment of hindbrain segmental identity requires signaling by FGF3 and FGF8. Curr Biol. 12, 1117–1123. (2002).

74. Hirsch, M., Glover, J., Dufour, H., Brunet, J. & Goridis, C. Forced expression of Phox2 homeodomain transcription factors induces a branchio-visceromotor axonal phenotype. Dev Biol. 303, 687–702. (2007).

75. Jarrar, W., Dias, J., Ericson, J., Arnold, H. & Holz, A. Nkx2.2 and Nkx2.9 are the key regulators to determine cell fate of branchial and visceral motor neurons in caudal hindbrain. PloS One 10, e0124408. (2015).

76. Andre, P., Song, H., Kim, W., Kispert, A. & Yang, Y. Wnt5a and Wnt11 regulate mammalian anterior-posterior axis elongation. Development 142, 1516–1527. (2015).

77. Nordström, U., Maier, E., Jessell, T. & Edlund, T. An early role for WNT signaling in specifying neural patterns of Cdx and Hox gene expression and motor neuron subtype identity. PLoS Biol. 4, e252. (2006).

78. van den Akker, E., et al. Cdx1 and Cdx2 have overlapping functions in anteroposterior patterning and posterior axis elongation. Development 129, 2181–2193. (2002).

79. Metzis, V. et al. Nervous System Regionalization Entails Axial Allocation before Neural Differentiation. Cell 175, 1105–1118.e1117. (2018).

80. Chang, J., Skromne, I. & Ho, R. CDX4 and retinoic acid interact to position the hindbrain-spinal cord transition. Dev Biol. 410, 178–189. (2016).

81. Skromne, I., Thorsen, D., Hale, M., Prince, V. & Ho, R. Repression of the hindbrain developmental program by Cdx factors is required for the specification of the vertebrate spinal cord. Development 134, 2147–2158. (2007).

82. Joshi, P., Darr, A. & Skromne, I. CDX4 regulates the progression of neural maturation in the spinal cord. Dev Biol. 449, 132–142. (2019).

83. Wymeersch, F., Wilson, V. & Tsakiridis, A. Understanding axial progenitor biology in vivo and in vitro. Development 148, dev180612. (2021).

84. Forlani, S., Lawson, K. & Deschamps, J. Acquisition of Hox codes during gastrulation and axial elongation in the mouse embryo. Development 130, 3807–3819. (2003).

85. Gouti, M. et al. A Gene Regulatory Network Balances Neural and Mesoderm Specification during Vertebrate Trunk Development. Dev Cell. 41, 243–261.e247. (2017).

86. Henrique, D., Abranches, E., Verrier, L. & Storey, K. Neuromesodermal progenitors and the making of the spinal cord. Development 142, 2864–2875. (2015).

87. Kondoh, H., Takada, S. & Takemoto, T. Axial level-dependent molecular and cellular mechanisms underlying the genesis of the embryonic neural plate. Dev Growth Differ. 58, 427–436. (2016).

88. Wymeersch, F. et al. Transcriptionally dynamic progenitor populations organised around a stable niche drive axial patterning. Development 146, dev168161. (2019).

89. Huangfu, D. & Anderson, K. Cilia and Hedgehog responsiveness in the mouse. Proc Natl Acad Sci U S A. 102, 11325–11330. (2005).

90. De Mori, R. et al. Joubert syndrome-derived induced pluripotent stem cells show altered neuronal differentiation in vitro. Cell Tissue Res. 396, 255–267. (2024).

91. Bangs, F., Schrode, N., Hadjantonakis, A.-K. & Anderson, K. Lineage specificity of primary cilia in the mouse embryo. Nat Cell Biol. 17, 113–122. (2015).

92. Soares, H., Carmona, B., Nolasco, S., Viseu Melo, L. & Gonçalves, J. Cilia Distal Domain: Diversity in Evolutionarily Conserved Structures. Cells 8, 160. (2019).

93. Wang, J. et al. Variable phenotypes and penetrance between and within different zebrafish ciliary transition zone mutants. Dis Model Mech. 15, dmm049568. (2022).

94. Chang, C. et al. Atoh1 Controls Primary Cilia Formation to Allow for SHH-Triggered Granule Neuron Progenitor Proliferation. Developmental Cell 48, 184–199. (2019).

95. Ong, T., Trivedi, N., Wakefield, R., Frase, S. & Solecki, D. Siah2 integrates mitogenic and extracellular matrix signals linking neuronal progenitor ciliogenesis with germinal zone occupancy. Nat Commun. 11, 5312. (2020).

96. Blaess, S. et al. Beta1-integrins are critical for cerebellar granule cell precursor proliferation. J Neurosci. 24, 3402–3412. (2004).

97. Akhshi, T. & Trimble, W. A non-canonical Hedgehog pathway initiates ciliogenesis and autophagy. J Cell Biol. 220, e202004179. (2021).

98. Zhang, K. et al. Primary cilia are WNT-transducing organelles whose biogenesis is controlled by a WNT-PP1 axis. 58, 139–154.e138. (2023).

99. Gouti, M. et al. In vitro generation of neuromesodermal progenitors reveals distinct roles for wnt signalling in the specification of spinal cord and paraxial mesoderm identity. PLoS Biol. 12, e1001937. (2014).

100. Lippmann, E. et al. Deterministic HOX patterning in human pluripotent stem cell-derived neuroectoderm. Stem Cell Reports 14, 632–644. (2015).

101. Afzal, Z. & Krumlauf, R. Transcriptional Regulation and Implications for Controlling Hox Gene Expression. J Dev Biol. 10, 4. (2022).

102. Wind, M. et al. Defining the signalling determinants of a posterior ventral spinal cord identity in human neuromesodermal progenitor derivatives. Development 148, dev194415. (2021).

103. Cooper, F. et al. Notch signalling influences cell fate decisions and HOX gene induction in axial progenitors. Development 151, dev202098. (2024).

104. Young, T. & Deschamps, J. Hox, Cdx, and anteroposterior patterning in the mouse embryo. J. Curr Top Dev Biol. 88, 235–255. (2009).

105. Takada, S. et al. Wnt-3a regulates somite and tailbud formation in the mouse embryo. Genes Dev. 8, 174–189. (1994).

106. Epstein, M., Pillemer, G., Yelin, R., Yisraeli, J. & Fainsod, A. Patterning of the embryo along the anterior-posterior axis: the role of the caudal genes. Development 124, 3805–3814. (1997).

107. Davidson, A. & Zon, L. The caudal-related homeobox genes cdx1a and cdx4 act redundantly to regulate hox gene expression and the formation of putative hematopoietic stem cells during zebrafish embryogenesis. Dev Biol. 292, 506–518. (2006).

108. Faas, L. & Isaacs, H. Overlapping functions of Cdx1, Cdx2, and Cdx4 in the development of the amphibian Xenopus tropicalis. Dev Dyn. 238, 835–852. (2009).

109. Amin, S. et al. Cdx and T Brachyury Co-activate Growth Signaling in the Embryonic Axial Progenitor Niche. Cell Rep. 17, 3165–3177. (2016).

110. Neijts, R., Amin, S., van Rooijen, C. & Deschamps, J. Cdx is crucial for the timing mechanism driving colinear Hox activation and defines a trunk segment in the Hox cluster topology. Dev Biol. 422, 146–154. (2017).

111. Mazzoni, E. et al. Saltatory remodeling of Hox chromatin in response to rostrocaudal patterning signals. Nature Neurosci. 16, 1191–1198. (2013).

112. Schyr, R., Shabtai, Y., Shashikant, C. & Fainsod, A. Cdx1 is essential for the initiation of HoxC8 expression during early embryogenesis. FASEB J. 26, 2674–2684. (2012).

113. Simons, M. et al. Inversin, the gene product mutated in nephronophthisis type II, functions as a molecular switch between Wnt signaling pathways. Nat Genet. 37, 537–543. (2005).

114. Veland, I. et al. Inversin/Nephrocystin-2 is required for fibroblast polarity and directional cell migration. PLoS One 8, e60193. (2013).

115. Lancaster, M., Schroth, J. & Gleeson, J. Subcellular spatial regulation of canonical Wnt signalling at the primary cilium. Nat Cell Biol. 13, 700–707. (2011).

116. Ocbina, P., Tuson, M. & Anderson, K. Primary cilia are not required for normal canonical Wnt signaling in the mouse embryo. PLoS One 4, e6839 (2009).

117. Vuong, L. & Mlodzik, M. The complex relationship of Wnt-signaling pathways and cilia. Curr Top Dev Biol. 155, 95–125. (2023).

118. Bishara, J., Keens, T. & Perez, I. The genetics of congenital central hypoventilation syndrome: clinical implications. Appl Clin Genet. 11, 135–144. (2018).

119. Dubreuil, V. et al. A human mutation in Phox2b causes lack of CO2 chemosensitivity, fatal central apnea, and specific loss of parafacial neurons. Proc Natl Acad Sci U S A. 105, 1067–1072. (2008).

120. Dubreuil, V. et al. Defective respiratory rhythmogenesis and loss of central chemosensitivity in Phox2b mutants targeting retrotrapezoid nucleus neurons. J Neurosci. 29, 14836–14846. (2009).

121. Li, A. & Nattie, E. CO2 dialysis in one chemoreceptor site, the RTN: stimulus intensity and sensitivity in the awake rat. Respir Physiol Neurobiol. 133, 11–22. (2002).

122. Kanbar, R., Stornetta, R., Cash, D., Lewis, S. & Guyenet, P. Photostimulation of Phox2b medullary neurons activates cardiorespiratory function in conscious rats. Am J Respir Crit Care Med. 182, 1184–1194. (2010).

123. Joubert, M., Eisenring, J., Robb, J. & Andermann, F. Familial agenesis of the cerebellar vermis. A syndrome of episodic hyperpnea, abnormal eye movements, ataxia, and retardation. Neurology 19, 813–825. (1969).

124. Assou, S. et al. Recurrent Genetic Abnormalities in Human Pluripotent Stem Cells: Definition and Routine Detection in Culture Supernatant by Targeted Droplet Digital PCR. Stem Cell Reports 14, 1–8. (2020).

125. Concordet, J. & Haeussler, M. CRISPOR: intuitive guide selection for CRISPR/Cas9 genome editing experiments and screens. Nucleic Acids Res. 26, W242–W245. (2018).

126. Doench, J. et al. Optimized sgRNA design to maximize activity and minimize off-target effects of CRISPR-Cas9. Nat Biotechnol. 34, 184–191. (2016).

127. Haeussler, M. et al. Evaluation of off-target and on-target scoring algorithms and integration into the guide RNA selection tool CRISPOR. Genome Biol. 17, 148. (2016).

128. Dobin, A. et al. STAR: ultrafast universal RNA-seq aligner. Bioinformatics. 29, 15–21. (2013).

129. Ewels, P., Magnusson, M., Lundin, S. & Käller, M. MultiQC: summarize analysis results for multiple tools and samples in a single report. Bioinformatics. 32, 3047–3048. (2016).

130. Liao, Y., Smyth, G. & Shi, W. featureCounts: an efficient general purpose program for assigning sequence reads to genomic features. Bioinformatics. 30, 923–930. (2014).

131. Lê, S., Josse, J. & Husson, F. FactoMineR: An R Package for Multivariate Analysis. Journal of Statistical Software 25, 1–18. (2008).

132. Love, M., Huber, W. & Anders, S. Moderated estimation of fold change and dispersion for RNA-seq data with DESeq2. Genome Biol. 15, 550. (2014).

133. Zink, R., Wolfinger, R. & Mann, G. Summarizing the incidence of adverse events using volcano plots and time intervals. Clin Trials. 10, 398–406. (2013).

134. Young, M., Wakefield, M., Smyth, G. & Oshlack, A. Gene ontology analysis for RNA-seq: accounting for selection bias. Genome Biol. 11, R14. (2010).

135. Alhamdoosh, M. et al. Combining multiple tools outperforms individual methods in gene set enrichment analyses. Bioinformatics. 33, 414–424. (2017).

